# Computed tomographic analysis of dental system of three Jurassic ceratopsians: implications for the evolution of the tooth replacement pattern and diet in early-diverging ceratopsians

**DOI:** 10.1101/2022.01.16.476516

**Authors:** Jinfeng Hu, Catherine A. Forster, Xing Xu, Qi Zhao, Yiming He, Fenglu Han

## Abstract

The dental system of ceratopsids is among the most specialized structure in Dinosauria, and includes high angled wear surfaces, split tooth roots, and multiple teeth in each tooth family. However, the early evolution of this unique dental system is generally poorly understood due to a lack of knowledge of the dental morphology and development in early-diverging ceratopsians. Here we study the dental system of three of the earliest-diverging Chinese ceratopsians: *Yinlong* and *Hualianceratops* from the early Late Jurassic of Xinjiang, and *Chaoyangsaurus* from the Late Jurassic of Liaoning. By using micro-computed tomographic analyses, our study has revealed significant new information regarding the dental system of these early ceratopsians, including no more than five replacement teeth in each jaw quadrant; at most one generation of replacement teeth in each alveolus; nearly full resorption of the functional tooth root during tooth replacement; and occlusion with low-angled, concave wear facets that differs significantly from the shearing occlusal system seen in ceratopsids. *Yinlong* displays an increase in the number of maxillary tooth alveoli and a decrease in the number of replacement teeth during ontogeny as well as the retention of remnants of functional teeth in the largest individual. Early-diverging ceratopsians thus display a relatively slow tooth replacement rate compared to late-diverging ceratopsians. Combined with paleobotany and palaeoenvironment data, *Yinlong* likely uses gastroliths to triturate foodstuffs, and the difference in diet strategy might have influenced the pattern of tooth replacement in later-diverging ceratopsians.

## Introduction

During the Cretaceous, the ceratopsids became one of the dominant herbivorous terrestrial clades and developed dental batteries composed of a large number of teeth that interlocked vertically and rostro-caudally in the jaw (Edmund, 1960; Dodson et al., 2004). Ceratopsids developed two-rooted teeth to facilitate vertical integration of the tooth batteries with up to four teeth in each vertical series (Edmund, 1960). This contrasts with non-ceratopsid taxa such as *Protoceratops* which retain single-rooted teeth which, although compacted rostro-caudally, have no more than two replacement teeth in each alveolus (Edmund, 1960). The early-diverging Early Cretaceous neoceratopsians, including *Auroraceratops* and *Archaeoceratops*, have only one replacement tooth in each alveolus (Tanoue et al., 2012). By using computed tomography, He et al. (2018) added more detailed information on the Early Cretaceous neoceratopsian *Liaoceratops* and presented evidence of the presence of the two replacement teeth per alveolus and shallow grooves on the roots to facilitate close-packing. Tracts of partially resorbed functional teeth in *Liaoceratops* appear to follow the growth of the jaws. *Liaoceratops* represents the earliest amniote for which multiple generations of tooth remnants are documented (He et al., 2018).

Here we investigate the tooth replacement pattern in even earlier-diverging Late Jurassic ceratopsians using micro-computed tomography (micro-CT) imaging. Three earliest-diverging ceratopsians were studied: *Yinlong downsi*, *Hualianceratops wucaiwanensis*, and *Chaoyangsaurus youngi* (Zhao et al., 1999; Xu et al., 2006; Han et al., 2015). *Yinlong* and *Hualianceratops* are from the upper Jurassic Shishugou Formation of Junggar Basin, Xinjiang, China (Xu et al., 2006; Han et al., 2015). *Yinlong* is one of the earliest and most complete ceratopsian dinosaurs and is known from dozens of individuals (Han et al., 2018), whereas *Hualianceratops* is known from only the holotype, a partial skull and mandible (Han et al., 2015). *Chaoyangsaurus* is from the Upper Jurassic Tuchengzi Formation of Liaoning Province, China, and is represented by a partial skull and paired mandibles (Zhao et al., 1999). This study provides crucial new evidence in our understanding of the initial evolution of ceratopsian dental specializations and diet.

## Materials & Methods

Three earliest-diverging ceratopsians *Yinlong*, *Chaoyangsaurus* and *Hualianceratops* were examined. The skull and dentary materials of *Yinlong* have been described in detail previously (Han et al., 2016; Han et al., 2018). Four skulls of *Yinlong downsi* are included in this study, IVPP V14530 (the holotype), IVPP V18636, IVPP V18637 and IVPP V18638 (Figs. 1-4).

**FIGURE 1.**
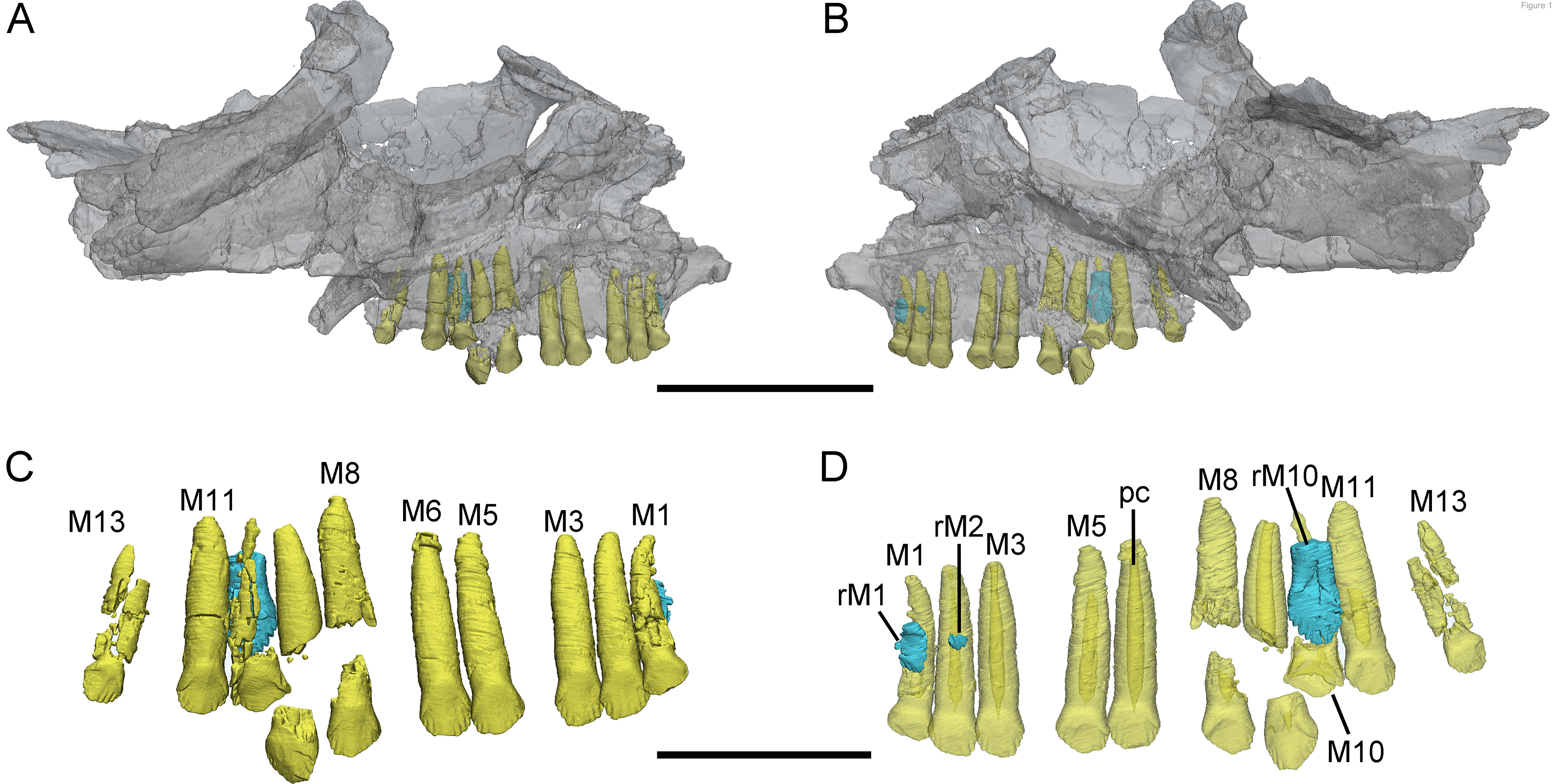
3D reconstructions of maxillary teeth in *Yinlong downsi* (IVPP V18638). Elements in the CT reconstructions are color-coded as follows: functional maxillary teeth, yellow; replacement teeth, cyan. (A) and (B), transparent reconstructions of the left maxilla of V18637 in labial and lingual view, respectively; (C), maxillary tooth row in labial view; (D), transparent reconstruction of the maxillary tooth row in lingual view. Abbreviations: M1-M13, the first to 13^th^ functional teeth in the maxilla; rM1, rM2, and rM10, the replacement teeth in first, third, and 10^th^ tooth alveolus; pc, pulp cavity. Scale bars equal 5 cm (A-B) and 2.5 cm (C-D).

**FIGURE 2.**
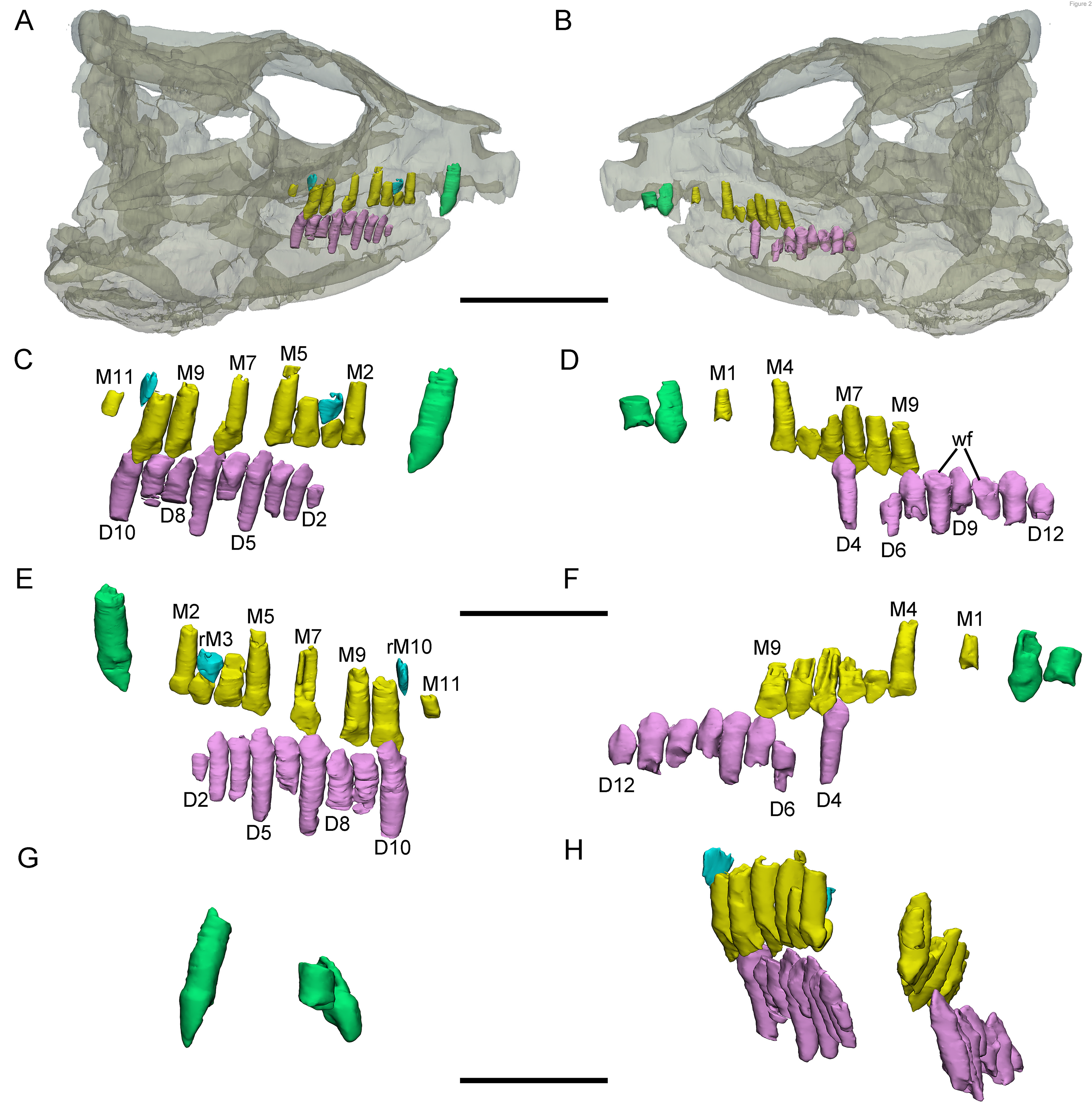
3D reconstructions of premaxillary and cheek teeth in *Yinlong downsi* (IVPP V18636). Elements in the CT reconstructions are color-coded as follows: functional premaxillary teeth, green; functional maxillary teeth, yellow; functional dentary teeth, lavender; replacement teeth, cyan. (A) and (B), transparent reconstruction of the skull of IVPP V18636 in right and left lateral view; (C) and (E), the right tooth rows in labial and lingual view; (D) and (F), the left tooth rows in labial and lingual view; (G), the premaxillary teeth in anterior view; (H), dentitions in the maxilla and dentary in anterior view. Abbreviations: M1-M11, first to 11^th^ functional teeth in the maxilla; rM3 and rM10, the replacement teeth in third and 10^th^ alveolus; D2-D12, second to 12^th^ functional teeth in the dentary; wf, wear surface. Scale bars equal 5 cm (A-B), 2.5 cm (C- F), and 2 cm (G-H).

**FIGURE 3.**
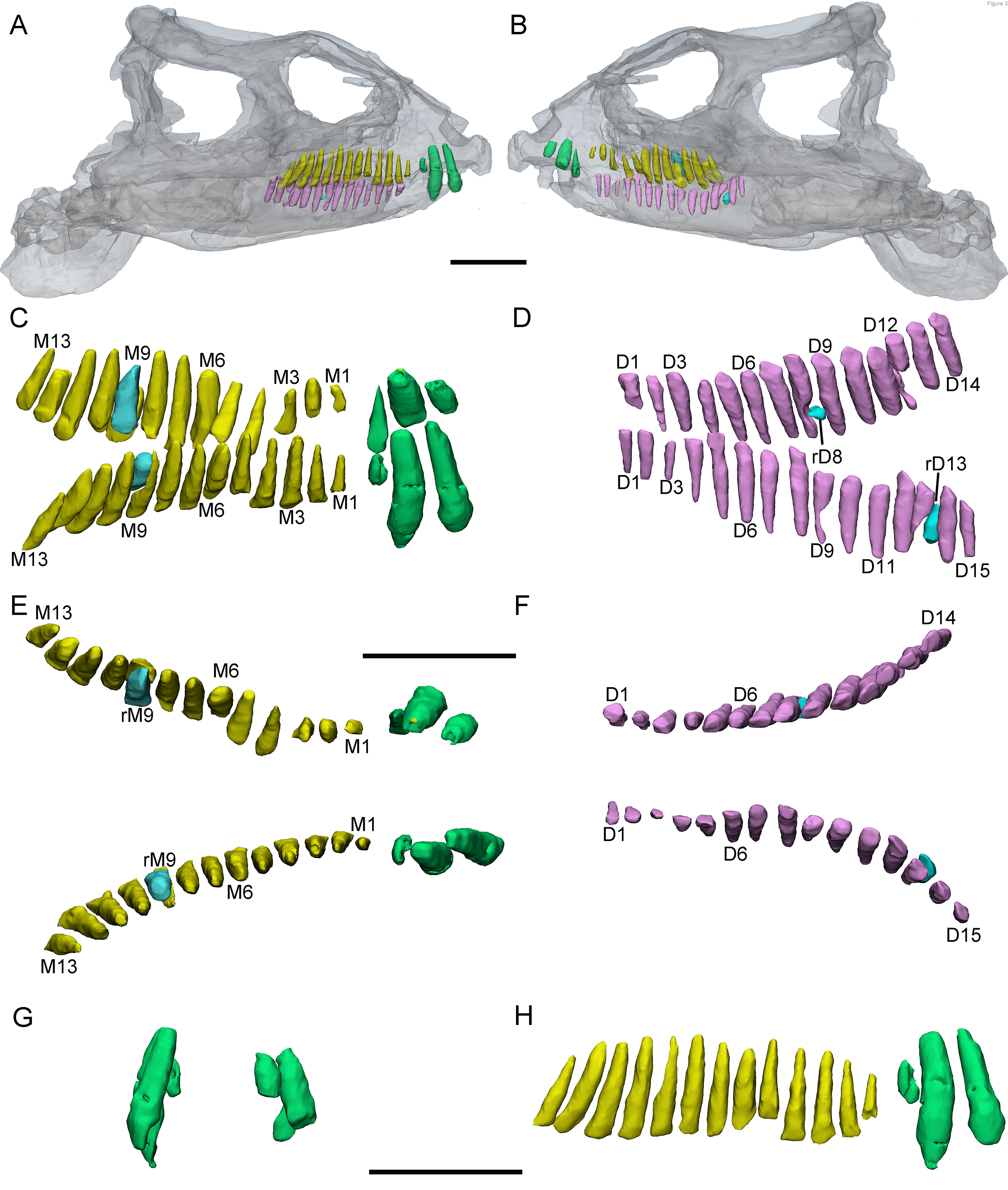
3D reconstructions of predentary and cheek teeth in *Yinlong downsi* (IVPP V14530). Elements in the CT reconstructions are color-coded as follows: functional premaxillary teeth, green; functional maxillary teeth, yellow; functional dentary teeth, lavender; replacement teeth, cyan. (A) and (B), transparent reconstructions of the holotype in right and left view; (C), reconstructions of the premaxillary and maxillary tooth rows in dorsal-labial view; (D), reconstructions of the dentary tooth rows in dorsal-labial view. (E) and (F), reconstructions of tooth rows in the upper and lower jaws in dorsal view; (G), reconstructions of the premaxillary teeth in anterior view; (H), reconstructions of the right tooth row in the upper jaw in labial view. Abbreviations: a, anterior; M1-M13, first to 13^th^ functional teeth in the maxilla; rM9, the replacement tooth in the ninth alveolus; D1-D15, second to 15^th^ functional teeth in the dentary; rD8 and rD13, the replacement teeth in the eighth and 13^th^ alveolus. Scale bars equal 5 cm (A-B) and 4 cm (C-H).

**FIGURE 4.**
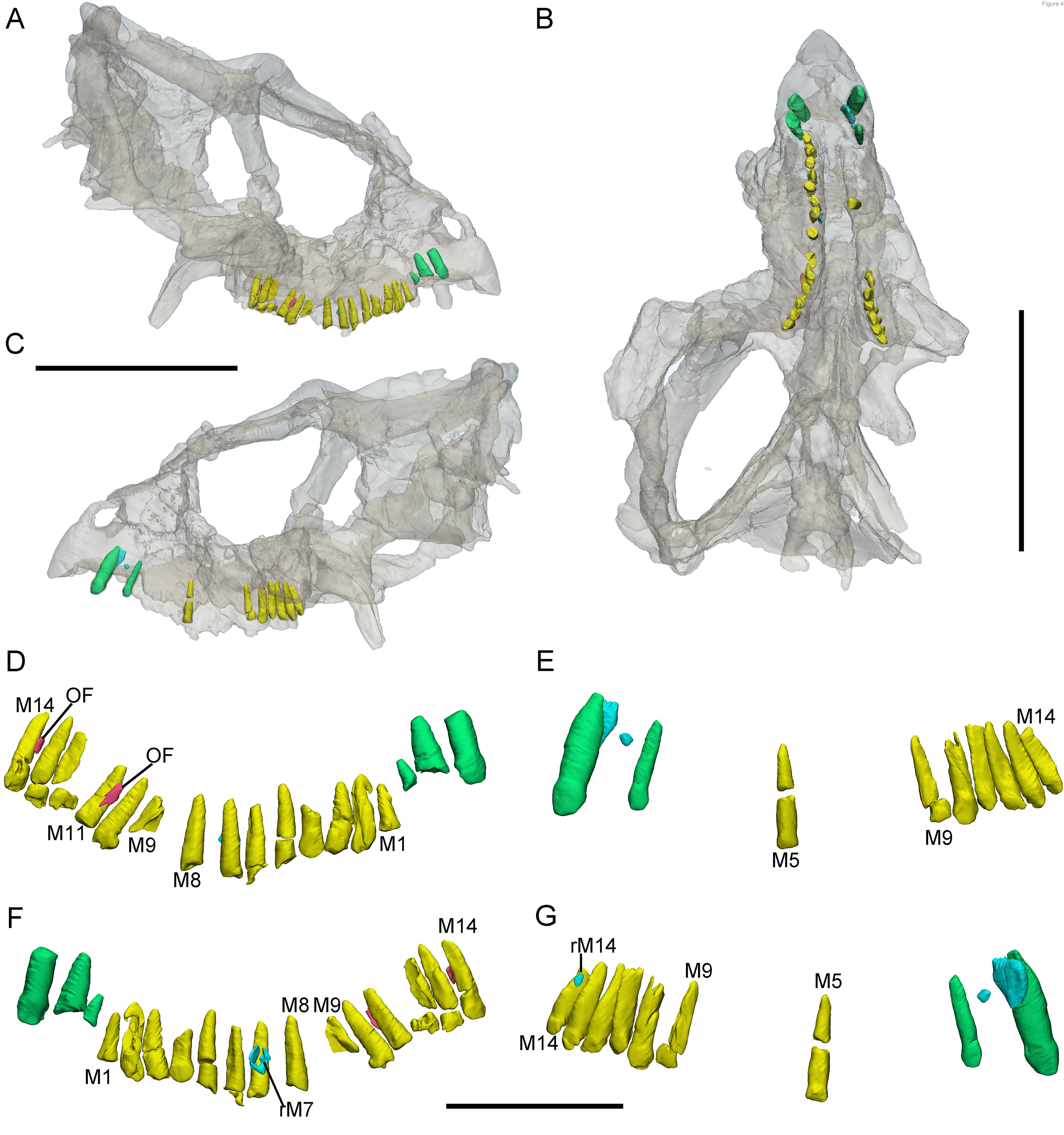
3D reconstructions of premaxillary and maxillary teeth in the largest specimen (IVPP V18637) of *Yinlong downsi*. Elements in the CT reconstructions are color-coded as follows: functional premaxillary teeth, green; functional maxillary teeth, yellow; replacement teeth, cyan; old functional teeth, red. (A) and (B), transparent reconstructions of the skull of V18637 in right and left view; (C), transparent reconstruction of the skull of IVPP V18637 in occlusal view; (D) and (E), two sides of tooth rows in the labial view; (F) and (G), two sides of tooth rows in the lingual view. Abbreviations: OF, old functional tooth; M1-M14, first to 14^th^ functional teeth in the maxilla; rM7, the replacement tooth in the seventh alveolus. Scale bars equal 10 cm (A, B, and C) and 4 cm (D-G).

Among them, IVPP V18637 is the largest specimen and IVPP V18638 is the smallest one. IVPP V18636 is slightly smaller than the holotype. The holotype of *Hualianceratops* (IVPP V18641) was also CT scanned, but its teeth were unidentified due to poor preservation. An additional specimen, IVPP V28614 (field number WCW-05A-2), which only preserves the left dentary is described here for comparison (Fig. 6). We assigned this specimen to *Hualianceratops* based on the deep and short dentary which measures 83.46 mm in length and has a depth of 33.38 mm at the rostral end (40% length) and strongly rugose sculpturing present on the lateral surface of the dentary (Fig. 6A) (Han et al., 2015). The holotype of *Chaoyangsaurus* (IGCAGS V371) includes the dorsal part of a skull and a nearly complete mandible (Fig. 5) (Zhao et al., 1999).

**FIGURE 5.**
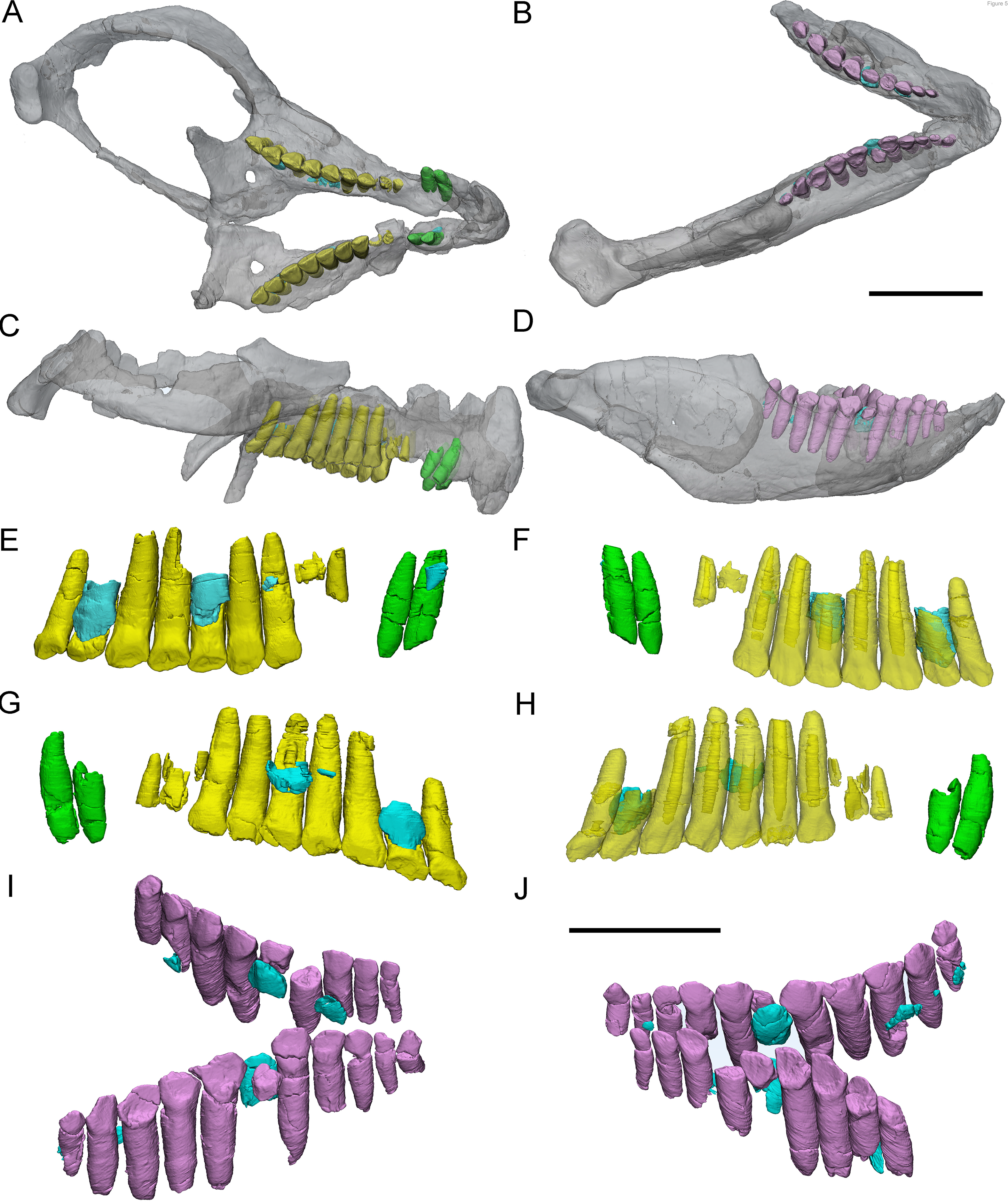
3D reconstructions of premaxillary and cheek teeth in *Chaoyangsaurus youngi* (IGCAGS V371). Elements in the CT reconstructions are color-coded as follows: functional premaxillary teeth, green; functional maxillary teeth, yellow; functional dentary teeth, lavender; replacement teeth, cyan. (A) and (B), transparent reconstructions of the maxilla and dentary in occlusal view; (C) and (D), transparent reconstructions of the maxilla and dentary in right lateral view; (E), reconstructions of the dentition in the left premaxilla and maxilla in lingual view; (F), transparent reconstructions of the dentition in the left premaxilla and maxilla in labial view; (G), reconstructions of the dentition in the right premaxilla and maxilla in lingual view; (H), transparent reconstructions of the dentition in the right premaxilla and maxilla in labial view; (I) and (J), reconstructions of the dentition in the dentary in right-dorsal and left-dorsal view. Scale bars equal 3 cm (A-D) and 2 cm (E-J).

**FIGURE 6.**
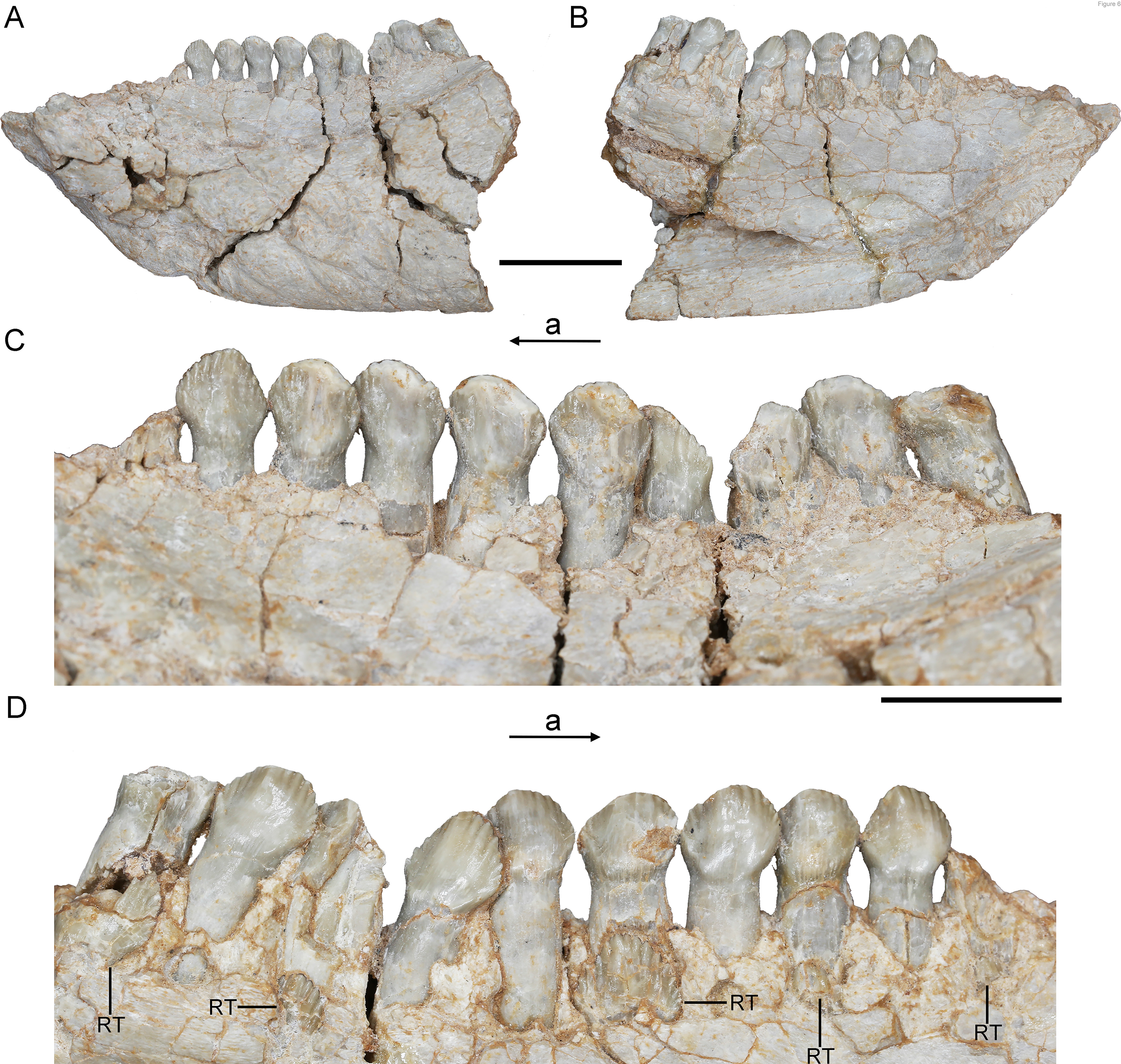
*Hualianceratops wucaiwanensis*, IVPP V28614 (WCW-05A-2). (A) and (B), the left dentary in labial and lingual view, respectively; (C) and (D), the tooth row of the dentary in the labial and lingual view, respectively. Abbreviations: a, anterior; RT, replacement tooth. Scale bars equal 2 cm (A-B) and 1 cm (C-D).

### Computed tomography

High-resolution X-ray micro-computed tomography was used to reveal internal anatomical features of tooth replacement in the maxilla and dentary. Scanning on IVPP V14530, IVPP V18636, and IVPP V18637 was carried out using a 450 kV micro-computed tomography instrument (450 ICT) at the Key Laboratory of Vertebrate Evolution and Human Origins.

Scanning on IVPP V18638 and IGCAGS V371 was carried out using a 300 kV micro-computed tomography instrument (Phoenix Vtomex M) and the detector (Dynamic41-100) at the Key Laboratory of Vertebrate Evolution and Human Origins. Scanning parameters of these specimens are displayed in TABLE 1.

**TABLE 1.**
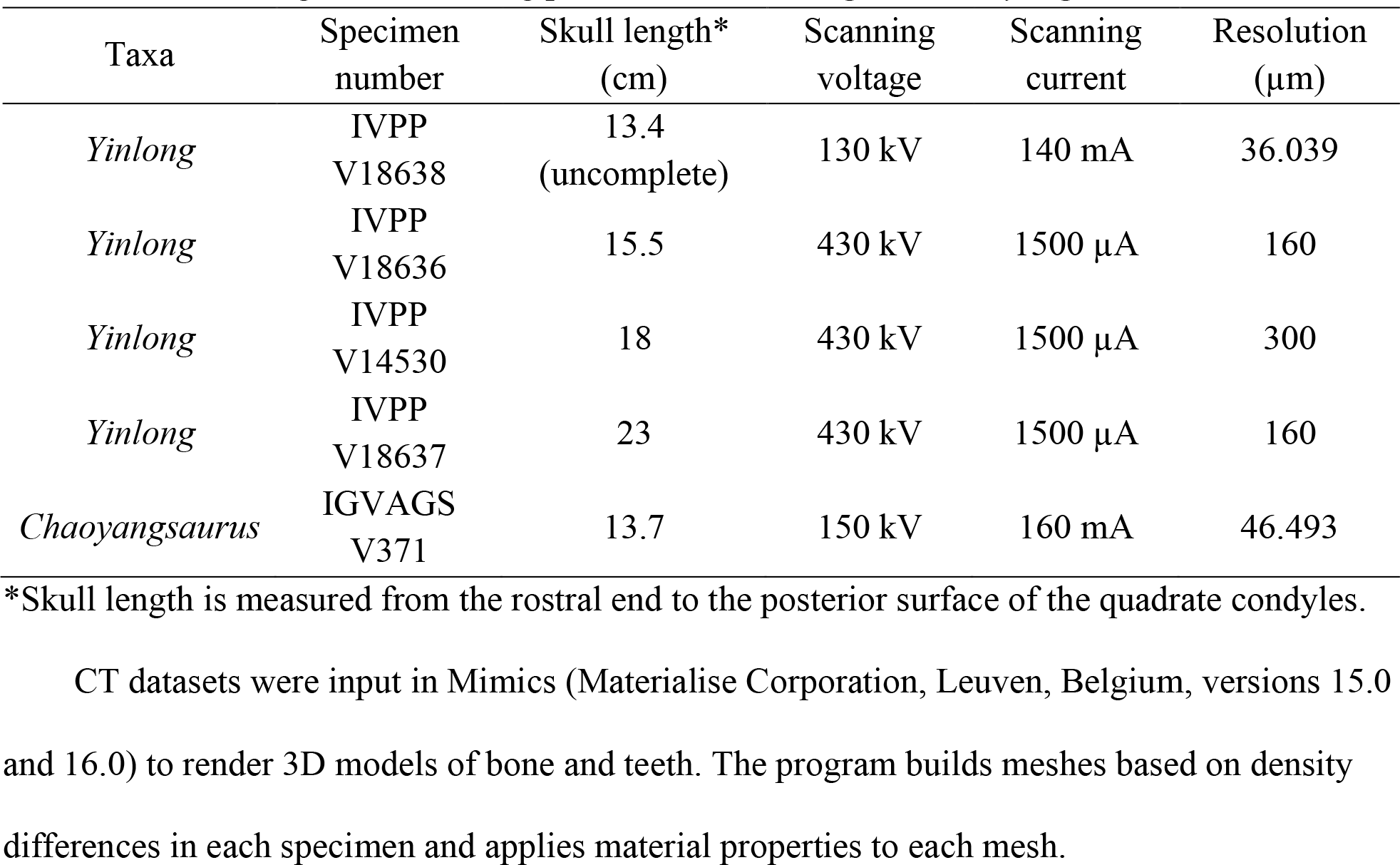
Skull length and scanning parameters of Yinlong and Chaoyangsaurus.

### The reconstruction of Zahnreihen

Edmund (1960) hypothesized that teeth in reptiles are replaced in an ordered, alternating segmented pattern called a Zahnreihe. Each Zahnreihe consists of a series of teeth, including unerupted teeth, where a more mesial tooth is more mature than a more distal one (Hanai & Tsuihiji, 2019). The distance between two successive Zahnreihe is the Z-spacing (Demar, 1972). Previous researches usually defined the Zahnreihen by measurements of teeth or by applying a replacement index (Demar & Bolt, 1981; Fastnacht, 2008; He et al., 2018). Because few replacement teeth are preserved in *Yinlong*, it is difficult to reconstruct the tooth replacement waves by applying the same replacement index used in *Liaoceratops*. Therefore, we reconstructed the Zahnreihen according to the degree of tooth wear and the location of replacement teeth, as used in *Shunosaurus* (Chatterjee & Zheng, 2002), as well as applying a new methodology that includes the developmental stage of the pulp cavity. We divided the functional teeth in *Yinlong* into four stages: (F1) no or slight wear on marginal denticles with an open pulp cavity; (F2) wear on marginal denticles and a slightly concave lingual wear facet with a large pulp cavity; (F3) extensive wear on marginal denticles and a concave lingual wear facet with the depression on the lingual surface of the roots or a bud of the replacement tooth; (F4) polished and greatly worn marginal denticles and a highly concave lingual wear facet with a broken pulp cavity or the emergence of a replacement tooth. Three stages of replacement teeth are recognized: (R1) small incipient tooth showing the tip of the crown; (R2) crown fully erupted; (R3) crown reaches the base of the functional crown.

Based on stage division, each functional tooth and replacement tooth was plotted on a graph whose vertical axis is the growth stage and horizontal axis is the tooth position. In the graph, these teeth show a regular pattern that the growth stage decreases progressively and periodically over several-tooth positions. Each degressive sequence represents a Zahnreihe indicated by a series of teeth linked with each other as black lines (Fig. 8). The distance between adjacent Zahnreihen is Z-spacing and the Z-spacing of *Yinlong* is described by the mean of all measurements.

## Results

### Dentition of the early-diverging ceratopsian *Yinlong*

IVPP V18638 only preserves the right maxilla (Fig. 1). In IVPP V18636, the premaxillae, the maxillae and the dentary are all well preserved, but some teeth are missing (Fig. 2). The holotype (IVPP V14530) preserves nearly complete maxillae and mandibles (Fig. 3), and the largest specimen (IVPP V18637) contains maxillae but lacks the mandibles (Fig. 4).

### Premaxillary teeth

All premaxillae bear three alveoli (Figs. 2-4), and all three teeth are preserved in IVPP V14530 (Fig. 3D). In IVPP V18636, the anterior two functional teeth are preserved in the left premaxilla and the second functional tooth is shown in the right premaxilla (Fig. 2A-2D). In the largest specimen (IVPP V18637), the second left functional premaxillary tooth has been lost and a replacement tooth remains in the alveolus (Fig. 4E and 4G). The right premaxilla is incomplete and the first tooth is poorly preserved but the second and third are present with minor damage to their roots (Fig. 4D and 4F).

The digital reconstructions show that the second functional premaxillary tooth is the largest among all maxillary or predentary teeth, and the third premaxillary tooth crown is quite short (Fig. 3A, 3C and 3H). The lateral surface of the premaxillary teeth is convex (Fig. 2G, 3E, 4B). Compared with the functional teeth on the maxilla, the long axes of the roots of the premaxillary teeth incline more dorsolingually (Fig. 2G and 2H, 3E, 4B).

All well-developed roots of all the functional teeth in the premaxilla are nearly conical and compressed labio-lingually into an oval cross-section. Compared with other premaxillary teeth, the tip and root of the second premaxillary tooth curves more posteriorly to appear arched in lateral view (Fig. 3C and 3H, 4D). The functional tooth crowns in the premaxilla are semiconical in shape and have similar rhomboidal outlines in lateral view. They taper apically without excessive wear (Fig. 2C, 3D and 3H, 4E and 4G). In rostral view, the crowns are slightly compressed labiolingually, with the lingual surface flattened and the labial rounded (Fig. 2G, 3G). The crown morphology of the first and second premaxillary tooth in IVPP V14530 is slightly different, with an abrupt step in the second premaxillary tooth between the inflated base and the more distal, lingually flattened crown (Fig. 3G) (Xu et al., 2006). The crown of the second tooth in IVPP V18636 also possesses this step but is more weakly developed than in IVPP V14530 (Fig. 2E and 2G).

Premaxillary replacement teeth are only preserved in the largest skull (IVPP V18637) (Fig. 4A and 4C). In lingual view, replacement teeth are present in the first and second alveoli of the left premaxilla (Fig. 4G). They are positioned lingual to their corresponding functional teeth although the functional tooth in the second alveolus is missing. The rootless replacement tooth in the first alveolus lies adjacent to the lingual wall of the functional tooth root. The apex of its crown is positioned halfway down the root of its functional tooth. (Fig. 4G). Slight resorption can be seen in the lingual side of the root of the left first functional tooth (Fig. 4E and 4G). The cross- section shows that the pulp cavity in the first replacement tooth is larger than that of the functional tooth, with a thinner layer of dentine. The apex of the first replacement tooth is more acuminate than that of the corresponding functional tooth (Fig. 4G). The first replacement tooth is nearly triangular in lingual and labial view with a mesiodistally elongated oval outline in cross- section and compressed labiolingually (Fig. 4G). The second replacement tooth in the left premaxilla is newly erupted and only preserves the tip of the crown. The replacement premaxillary teeth in *Liaoceratops* have cone-shaped crowns and are similar in morphology to their corresponding functional teeth (He et al., 2018). In *Liaoceratops*, 1-2 replacement teeth exist in each premaxillary alveolus.

### Maxillary teeth

The incomplete right maxilla of IVPP V18638 contains 10 functional teeth and 3 empty alveoli (Fig. 1A). The left and right maxillae of IVPP V18636 contain 7 functional teeth and 8 functional teeth respectively with some empty sockets (Fig. 2C and 2D). According to cross-sections, four empty sockets in the left maxilla and three empty sockets in the right maxilla can be discerned in IVPP V18636 (Table. 2). Both the left and right maxillae of the holotype contain 13 functional teeth as identified before (Fig. 3C and 3E) (Xu et al., 2006; Han et al., 2016). However, in the largest specimen IVPP V18637, the incomplete maxillae contain 7 functional teeth and 14 functional teeth on the left and right sides respectively (Fig. 4D and 4E). The left maxilla of IVPP V18637 contains seven empty sockets, suggesting that the maxilla bears 14 or more teeth in adult *Yinlong*.

**TABLE 2.**
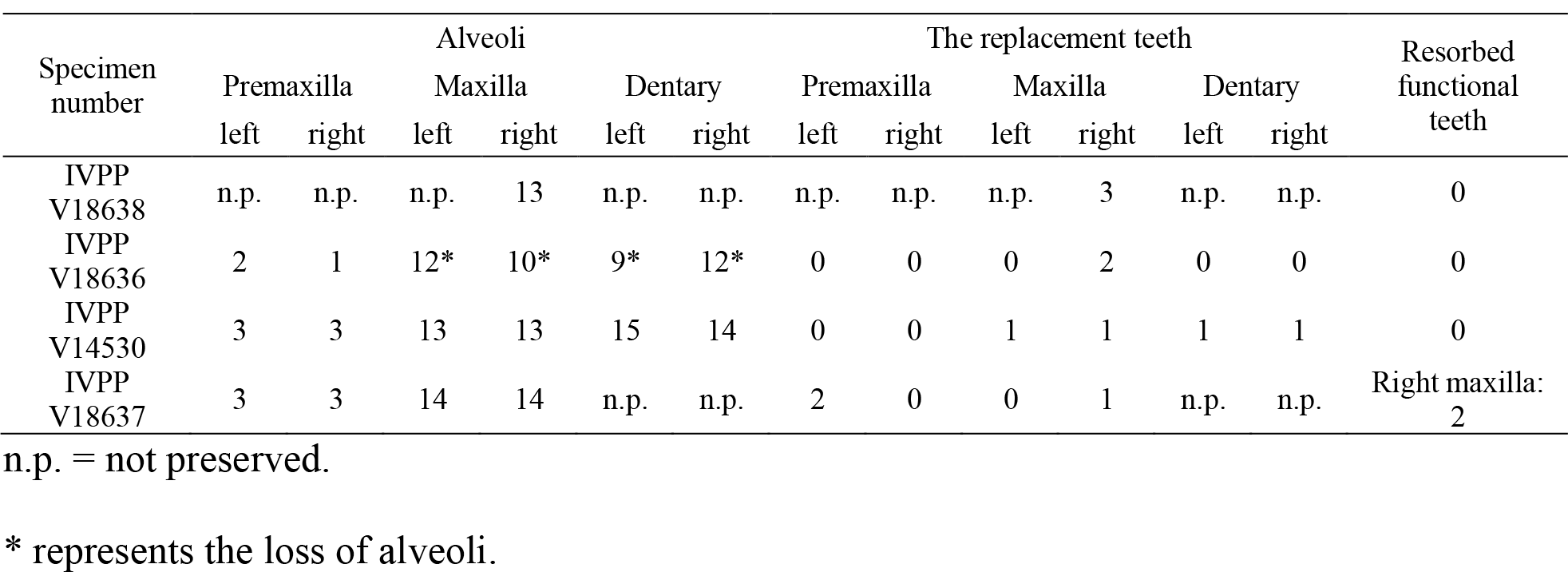
List of the ontogenetic difference in specimens of Yinlong.

The maxillary tooth row is curved inwards (Fig. 3E, 4B). Generally, the length of functional teeth increases to a maximum in the middle part of the maxillary tooth row and then decreases caudally (Figs. 1-4). All roots of functional teeth are widest at their crown bases and taper apically to form elongate roots with a subcircular cross-section (Fig. 1C, 3H, 4D). The root cross- sections reveal a pulp cavity surrounded by a thick layer of dentine. According to our 3D reconstructions and cross-section, the pulp cavities of some functional teeth are open at their tips such as M3 and M9 in IVPP V18638 and the functional teeth with the open pulp cavity have a thinner layer of dentine (Fig. 1D). The elongate pulp cavity in the functional tooth nearly extends over the whole root (Fig. 1D). In all specimens, strong root resorption is seen on the lingual surface of some functional teeth adjacent to replacement teeth (Fig, 1D, 2F, 3C and 3D). In these cases, the dentine has been resorbed by the replacement teeth such that the root base has been hollowed (Fig. 3C). The root of M4 on the right maxilla of the holotype is also hollowed, but no replacement tooth is present on both sides (Fig. 3C). M4s are hollowed less than D8 which is attached by a replacement crown tip. Therefore, M4 may represent the primary stage of the resorption progress prior to replacement tooth development.

The crowns of functional teeth in the maxilla have a spatulate outline in labial view and are slightly bulbous at the base (Fig. 1C, 2C and 2D, 3H, 4E and 4F). In IVPP V18638 all of the crowns are relatively complete with the apex of most of the crowns worn slightly except M1 and M10 (Fig. 1C). The mesiodistal length and labiolingual width of erupted crowns increase to a maximum at their base. In labial view, several denticles are distributed over the margin beneath the base of the crown (Fig. 1C). About four denticles are distributed over the mesial and distal carinae of tooth crowns respectively and all the denticles are subequal in size and taper apically (Fig. 1C). This feature is present but weak in *Chaoyangsaurus*, *Psittacosaurus*, *Liaoceratops* and *Archaeoceratops* (Tanoue et al., 2009; Ryan et al., 2010). The primary ridge is prominent in M13 of V18638 and positioned in the midline of the crown. The lingual surface of crowns is concave except for M9 whose lingual surface is convex (Fig. 1D). In addition, M10, which is in the replacement process, has a more concave lingual surface than other functional teeth that have not undergone resorption. Therefore, we hypothesize that the lingual surfaces of the crowns are flat and gradually become concave as the wear facet develops (Fig. 1D). Similar wear facets can be seen in *Heterodontosaurus tucki* (Sereno, 2012).

The count of the replacement teeth in the maxilla of *Yinlong* is 1 out of 13 functional teeth in the holotype. The smallest specimen IVPP V18638 has the most replacement teeth in the maxilla with and CT data reveal 3 replacement teeth out of 10 functional teeth inside the right maxilla (Fig. 1D). The replacement (rM10) occurs lingual to M10 whose root has been almost completely resorbed with only a fragmentized layer of dentine remaining. This replacement tooth is well developed and consists of the complete crown and partial root. The apex of rM10 meets the base of the crown of the functional tooth. Compared with the functional teeth, the crowns of the replacement teeth are rhomboidal in labial view, compressed labiolingually, and the denticles extend nearly the entire margin of the crown (Fig. 1D). In IVPP V18636, there are two replacement teeth preserved in the right maxilla (Fig. 2C and 2E). The first replacement tooth, preserving only the crown, is attached to the dorsal side of M3. The base of the corresponding functional tooth has been hollowed and the root has been resorbed although the crown is still functional (Fig. 2D). In IVPP V18636, the crown of rM10 (rM10?) is positioned posterior to M10 and is similar to the premaxillary replacement tooth of V18637in having a triangular outline in lingual view (Fig. 4G). It may suggest that the replacement tooth with a compressed shape is relatively common in *Yinlong*.

Remnants of roots of resorbed functional teeth occur in IVPP V18637. The remnants are positioned labiodistal to functional M11 and M14 in the right maxilla (Fig. 4A and 4D).

Remnants of resorbed functional teeth preserve a thin layer of dentine and exhibit a crescent outline in cross-section. There is only one generation of resorbed tooth remnants along the maxillary tooth row. Remnants of resorbed functional teeth are also reported in *Liaoceratops*, *Coelophysis,* and a hadrosaurid, but the number of resorbed functional teeth in *Liaoceratops* are far more than *Yinlong*.(Bramble et al., 2017; Leblanc et al., 2017; He et al., 2018). In the holotype of *Liaoceratops*, about 28 remnants of the functional teeth are preserved in the right maxilla and at most four generations of teeth remnants are located at the middle part of the tooth row.

### Dentary teeth

The holotype has a complete dentary containing 15 functional teeth on the left and 14 functional teeth on the right (Fig. 3D and 3F). The dentaries of IVPP V18636 are incomplete, containing 9 functional teeth and 1 empty socket on the right dentary and 8 functional teeth and 3 empty sockets on the left (Fig. 2C and 2D). The left dentary is distorted so that the long axes of functional teeth in the two sides extend in different directions (Fig. 3H). The size of the dentary teeth increases to a maximum on the tooth 5 and 6 and then decreases. In dorsal view, the functional teeth in the middle of the dentary tooth row are compressed and their long axes incline ventromedially (Fig. 3F).

The morphologies of dentary roots are similar to the maxillary teeth with a nearly conical shape and oval cross-sections (Fig. 2E and 2F, 3D). Most functional teeth in the dentary have complete crowns (Fig. 2C and 2D). In labial view, the outline of functional teeth on the dentary is similar to maxillary teeth but its labial surface is concave (Fig. 2D). This concave surface has never been found in other ceratopsians, suggesting that *Yinlong* had relatively precise occlusion. The roots of the dentary teeth are similar to those of the maxillary teeth.

Two replacement teeth can be seen in the dentary of the holotype (Fig. 3D). Among them, the crowns of D9 in the left dentary and D12 in the right dentary have been hollowed although no replacement tooth is preserved. However, the cavity caused by the resorption is similar to its corresponding functional tooth on the contralateral side (Fig. 3D and 3F). It can be concluded that they are at a similar stage of the replacement progress. In addition, D8 on the right side also exists a replacement tooth. Patterns of symmetry in replacement patterns can be seen in *Yinlong*, but the replacement stage between two dentaries is slightly different.

### Dentition of Chaoyangsaurus

The holotype of *Chaoyangsaurus* (IGCAGS V371) preserves the last two premaxillary teeth. The premaxillary teeth of *Chaoyangsaurus* are ellipsoidal in cross-section and the crowns are not expanded mesiodistally at their base as in *Yinlong.* The apices of the crowns are missing (Fig. 5E-5H). The first preserved functional tooth in the left premaxilla is undergoing replacement and its corresponding replacement tooth crown is triangular with the apex inclined mesially (Fig. 5E). The long axis of the replacement tooth in the premaxilla retains the same angle of tilt with its corresponding functional tooth (Fig. 5E).

CT reconstructions reveal that the maxillary teeth of *Chaoyangsaurus* possess different crown morphology from *Yinlong*. In *Chaoyangsaurus*, the primary ridges are located more distally (Fig. 5E and 5G) and the basal ridge extends over more than 70% of the crown with denticles spread over the mesial and distal margins (Fig. 5F). The lingual surfaces of the maxillary crowns are concave and the crowns in the dentary also show concave surfaces similar to the situation in *Yinlong* (Fig. 5E, 5I and 5J). The concave surface in the lingual side of maxillary crowns and the labial side of dentary crowns may indicate wear facets similar to those of *Yinlong*. The roots of the teeth in *Chaoyangsaurus* are elongated and inclined lingually. CT data also reveals the phenomenon that the fourth and seventh functional teeth have pulp cavities open at their top and these teeth show less wear than others (Fig. 5F and 5H). Therefore, the functional teeth with open pulp cavities may be newly erupted. There are three replacement teeth out of nine functional teeth on both maxillary tooth rows (Fig. 5E and 5G).

The morphology of the dentary teeth is similar to that of maxillary teeth although no primary ridge or denticles exist on the dentary crowns (Fig. 5I and 5J). The left dentary of *Chaoyangsaurus* possesses three replacement teeth out of nine functional teeth and on the other side there are five replacement teeth out of 11 functional teeth (Fig. 5I and 5J). According to 3D reconstructions of maxillary and dentary teeth, the pulp cavity is gradually reduced through time after tooth eruption (Fig. 5F and 5H). The number of replacement teeth in *Chaoyangsaurus* is slightly more than that of *Yinlong*.

### Dentition of Hualianceratops

The crowns of the teeth in *Hualianceratops* are similar to those of *Yinlong*, but include more denticles along the margins (Fig. 6C and 6D). The dentary preserves the complete morphology of the crowns. They are subtriangular in labial view and the mesial and distal margins bear about seven denticles respectively (Fig. 6C). Ten functional dentary teeth are identified. The tooth crowns are slightly imbricated with the distal margin of each tooth overlapping the lingual side of the mesial margin of the preceding tooth. The first functional tooth is broken with only part of the root remains. Five replacement teeth are exposed on the lingual aspect of their corresponding functional teeth and exposed at the border of the alveoli (Fig. 6D).

### Replacement progress and tooth development in *Yinlong* and *Chaoyangsaurus*

In *Yinlong* and *Chaoyangsaurus*, the resorption of the functional tooth is initiated before the successional tooth has germinated (Fig. 1D, 5G and 5I). The functional tooth roots are resorbed resulting in a depression on the middle part of the roots (Fig. 1D and 5I). After the depression extends enough, the replacement teeth form lingual to the functional tooth roots with the crown situated a small distance away from the middle part of the roots. The replacement tooth crown then gradually grows out towards the margin of the alveolus. The most immature replacement teeth are represented by small cusps (Fig. 1C, 5E and 5G). With ontogeny, the crowns of more mature teeth become fully developed and largely resorb the lingual aspects of the roots of the functional teeth, part of which are housed in their pulp cavities (Fig. 1C, 3D, 4G and 5G). In this stage, some crowns in *Yinlong* and *Hualianceratops* of replacement teeth were flat labiolingually and possibly kept this morphology until erupted (Fig. 1D, 2C, 4G and 6D). However, the replacement crowns in *Chaoyangsaurus* were inflated and the morphology was almost the same as that of the functional teeth (Fig. 5E and 5G). Differing from the maxillary teeth, the crowns of the premaxillary replacement teeth are housed in the more apical part of the functional tooth and a similar situation occurs in *Chaoyangsaurus* (Fig. 4E, 4G and 5E). As the lingual surface of the functional teeth becomes heavily resorbed, the replacement teeth reach about 60% or more of their predicted full size (Fig. 1D). When the replacement tooth grows to its final size, most of the roots of the predecessors have faded through heavy resorption and may leave the small root remnants on the labial surface of its successor’s tooth (Fig. 7A and 7B, 7D and 7E).

**FIGURE 7.**
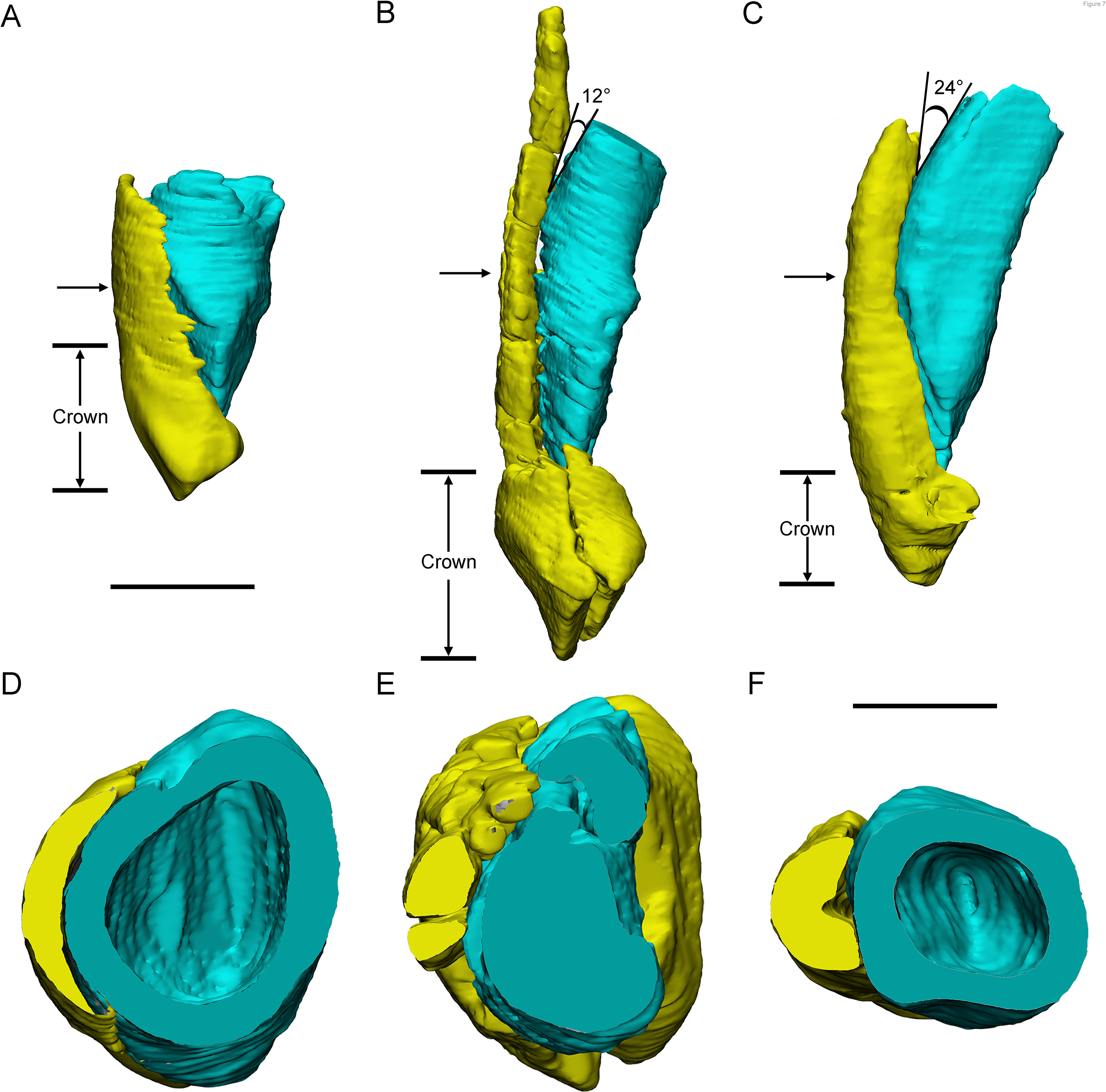
Three different replacement processes illustrated by teeth at similar replacement stage of *Chaoyangsaurus* (A and D), *Yinlong* (B and E), and *Liaoceratops* (C and F). Elements in the figure are color-coded as follows: functional maxillary teeth, yellow; replacement teeth, cyan. (A), the reconstruction of the tooth eight in the left maxilla of IGCAGS V371 in distal view; (B), the reconstruction of the tooth 10 in mesial view in IVPP V18638; (C), the reconstruction of the tooth seven in the right maxilla of the holotype of *Liaoceratops* (IVPP V12738) in mesial view; (D), (E) and (F), the cross-sections through the base of the replacement teeth of A, B, and C. The arrows of A, B, and C indicate where the cross-sections generate. The replacement teeth here have developed the complete crown and part of the root. The root of the replacement tooth in *Liaoceratops* inclines lingually and that in *Yinlong* also inclines lingually but with a smaller angle of inclination. The root of the replacement tooth in *Chaoyangsaurus* clings to its corresponding functional tooth tightly. The resorbed area on the functional tooth is larger in *Chaoyangsaurus* and *Yinlong* than in *Liaoceratops* because of the larger contact area. Therefore, the resorption degree of the functional tooth in *Chaoyangsaurus* and *Yinlong* is also larger than *Liaoceratops*. Scale bars equal 5 mm (A-C) and 2.5 mm (D-F).

### The Zahnreihen in *Yinlong*

In the Zahnreihen graph of IVPP V18638, these teeth show the regular pattern that the growth stage decreases progressively over a two-tooth position or three-tooth position period and hence at least four Zahnreihe are possibly identified (Fig. 8). The resulting Zahnreihen are formed by M1 to M3, M5 to M6, M8 to rM10 and M10 to M11 respectively and run more or less parallel to each other (Fig. 8). The M1-M3 and M8-M10 are well-defined tooth replacement series and the exceptions are rM1, rM2, and M13. In *Yinlong*, Z-spacing is between 2.5 and 3.0, and the average Z-spacing is 2.83. Edmund (1960) suggested that the Z-spacing in reptilian dentitions is higher in the anterior region of the tooth row generally. This pattern is also present in *Yinlong*, whereas Z-spacing is higher in the posterior region of the tooth row in *Liaoceratops* (He et al., 2018). Fastnacht (2008) suggested that the replacement ratio of tooth formation against tooth resorption can be directly derived by the Z-spacing. Although the replacement ratio cannot yield the time information, it represents the replacement rate to a certain extent and is only comparable within a single taxon. The lower the value is, the higher is the tooth replacement rate (Fastnacht, 2008). Therefore, Z-spacing provides an index to compare the replacement rate in one taxon or jaw element.

**FIGURE 8.**
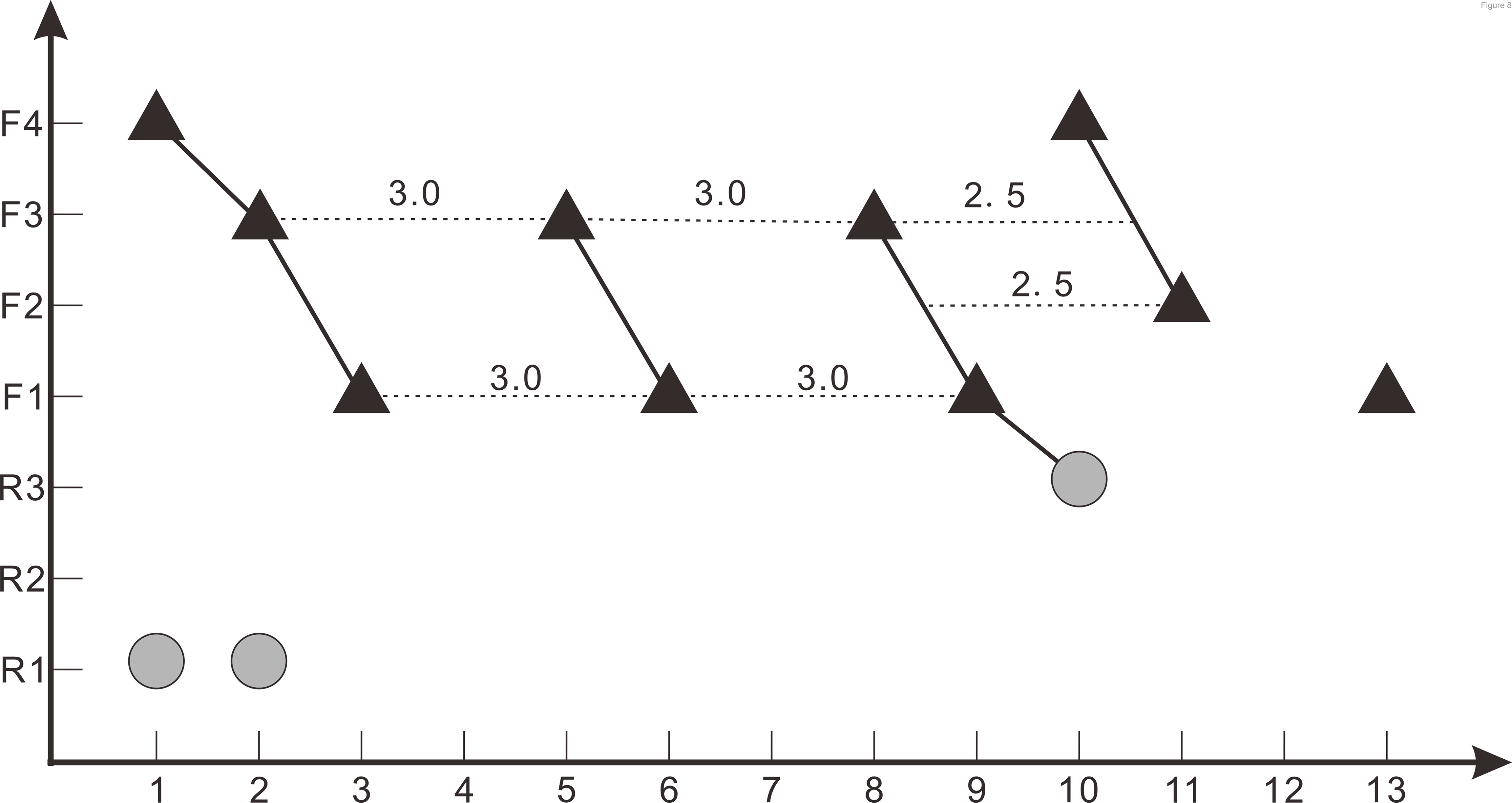
Z-spacing diagram of *Yinlong downsi* (IVPP V18638). The X-axis is the tooth position, Y-axis is the tooth replacement stage. The black triangle represents the functional tooth and the gray circle represents the replacement tooth. Four Zahnreihen are identified in this diagram marked as a series of teeth linked with each other as black lines. Each red imaginary line represents the Z-spacing which is the distance between Zahnreihen whose unit is a tooth position.

The lower Z-spacing in the posterior region of *Yinlong* may suggest that the posterior region of the tooth row has a higher replacement rate. To maintain the efficiency of chewing, it is advantageous to replace more worn teeth more rapidly. Therefore, this may indicate that the posterior region of the jaw in *Yinlong* is used more than the rostral portion to chew food. The situation in *Liaoceratops* is opposite that in *Yinlong* in that the anterior jaw region has the higher replacement rate and the food preparation may therefore occur more frequently in the anterior region (He et al., 2018). It is still not clear whether the transition of the place in the jaw the preparation of food mainly occurs is a universal phenomenon during the evolution of ceratopsians or it only happens in early-diverging taxa.

Demar (1972) reported that the value of the Z-spacing ranges from 1.56 to 2.80 in most reptiles. Z-spacing as the quantitative index could be used to assess the replacement patterns and avoid arbitrary interpretation of replacement patterns and facilitates objective comparison of patterns between different jaw elements, individuals, growth stages, taxa and so forth (Hanai & Tsuihiji, 2019). In *Liaoceratops*, the spacing between Zahnreihe ranges from 2.16 to 2.90 with a mean value of 2.58 (He et al., 2018). So far, only the Z-spacings of *Yinlong* and *Liaoceratops* are known in ceratopsians and more researches on the Z-spacing of ceratopsians are required to make meaningful comparisons. In non-avian dinosaurs, all known Z-spacing values are greater than 2.0 (Chatterjee & Zheng, 2002; Weishampel et al., 2004; Wiersma & Sander, 2017; Hanai & Tsuihiji, 2019; Becerra et al., 2020). Hanai & Tsuihiji (2019) examined some extant crocodiles such as *Alligator mississippiensis* and *Crocodylus siamensis* which present infrequent Z-spacing less than 2.0. These values indicate the replacement wave direction which is mesial-to-distal when Z-spacing is greater than 2.0, reversed when less than 2.0 and replaced in simple alternation between odd- and even-numbered tooth positions when exactly 2.0 (Hanai & Tsuihiji, 2019).

This indicates that new teeth erupt from distal to mesial order in either odd- and even-numbered alveoli in the maxilla of *Yinlong*.

## Discussion

### Ontogenetic changes in dentitions of *Yinlong*

Although the accurate ontogenetic stage of these four specimens is not clear, the ontogenetic variation of the tooth replacement pattern in this taxon can be discussed relative to their size difference. Previous research suggests that the maxilla in *Yinlong downsi* bears 13 teeth (Han et al., 2016). Our 3D reconstructions reveal that 13 functional alveoli are preserved in the maxilla of V18638 and a larger individual (IVPP V14530). However, the count of functional teeth in the largest individual (IVPP V18637) is at least 14. Hence the number of the maxillary teeth may increase with the ontogeny of *Yinlong downsi*, as in *Psittacosaurus mongoliensis* and *Protoceratops* (Brown & Schlaikjer, 1940; Sereno, 2006; Czepiński, 2020). In large individuals (IVPP V18637, IVPP V14530), there is one replacement tooth out of 14 or 13 functional teeth in the maxilla whereas smaller specimens (IVPP V18637, IVPP V18638) have a higher ratio of the replacement teeth to the functional teeth such as two RT/8 FT and three RT/10 FT (TABLE. 2).

This phenomenon may reflect that the early ontogenetic stage specimens of *Yinlong* may have a faster tooth replacement rate. As noted by He et al. (2018), remnants of mostly resorbed functional teeth are present in both juvenile and adult specimens of *Liaoceratops*. But the remnants of resorbed functional teeth are only present in the largest specimen (IVPP V18637) of *Yinlong*. Therefore, we conclude that the resorption rate may decrease in the ontogeny of *Yinlong*.

### The evolution of dental anatomy and replacement pattern in Ceratopsia

#### Dental anatomy

Tanoue et al. (2009) have presented that the evolutionary trend in dentitions of early-diverging ceratopsians includes an increase in the angle of the wear facets, development of a prominent primary ridge, development of deep indentations on the mesial and distal sides of the primary ridge and increase in size in neoceratopsians. By computed tomographic analysis, we found that the dentitions in *Yinlong*, *Hualianceratops* and *Chaoyangsaurus* exhibit new features compared to neoceratopsians including small numbers of teeth in tooth rows, concave surfaces on the lingual side of the maxillary crowns and labial side of the dentary crowns, loosely packed tooth rows, and regular occlusal surfaces. There are also some differences between these taxa.

The crowns of unworn teeth in *Yinlong* and *Hualianceratops* are subtriangular and bear primary ridges located at the midline of the crowns (Fig. 1C and 6C). Unlike *Yinlong* and *Hualianceratops*, the maxillary dentitions of *Chaoyangsaurus* developed ovate crowns and the relatively prominent primary ridge located at relatively distal to the midline of the crowns as in most neoceratopsians (Fig. 5F and 5H). In addition, the roots in *Yinlong* are straight, unlike *Chaoyangsaurus* whose functional roots are curved lingually (Fig. 9A-C). Overall, the dentitions of *Yinlong* and *Hualianceratops* have primitive conditions compared to *Chaoyangsaurus*.

**FIGURE 9.**
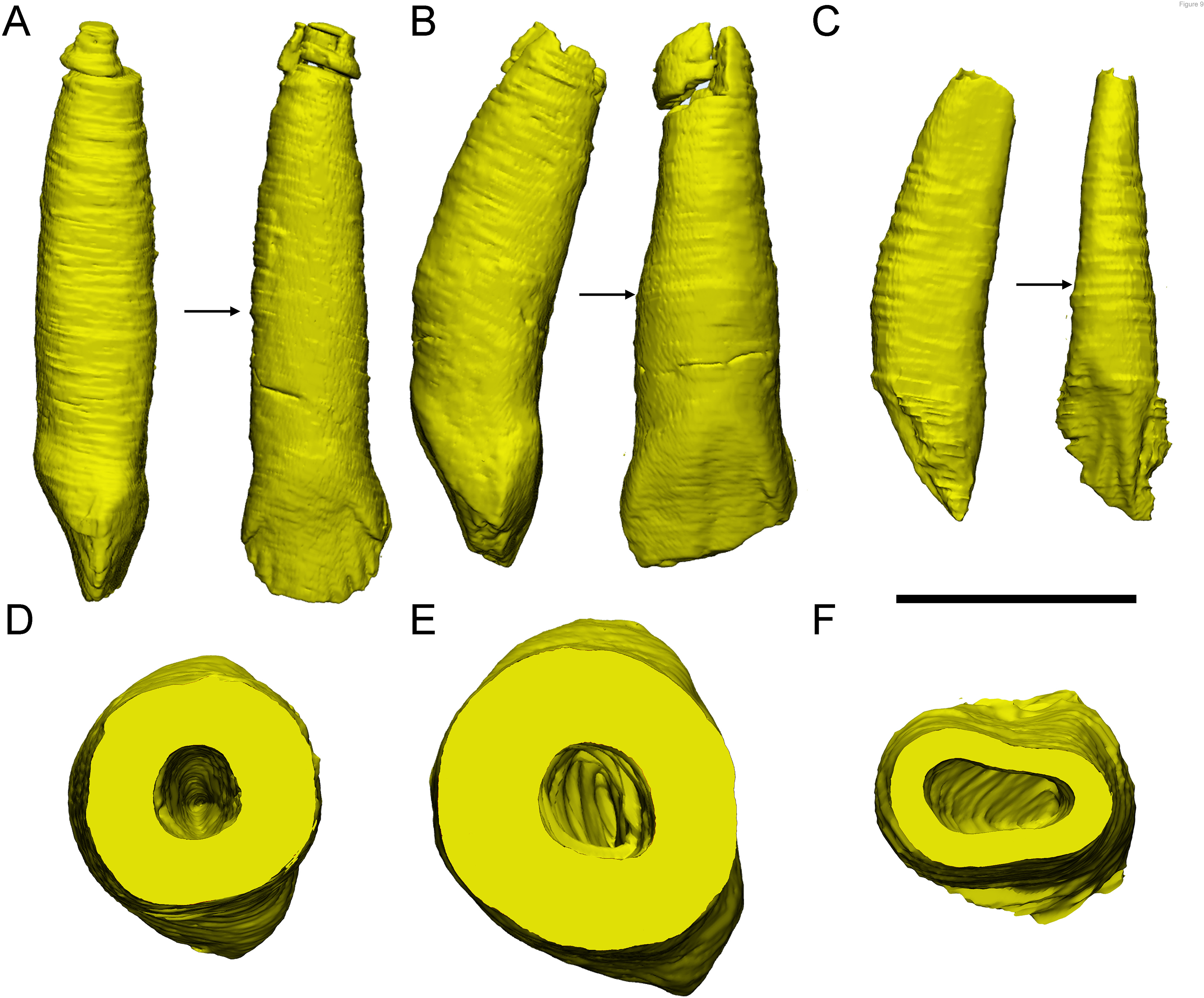
The reconstructions of three functional maxillary teeth at the middle part of the tooth row. (A), the reconstruction of tooth 6 in *Yinlong* (IVPP V18638) in mesial and labial view; (B), the reconstruction of tooth 7 in the left maxilla of IGCAGS V371 (*Chaoyangsaurus*) in mesial and distal view; (C), the reconstruction of tooth 9 in the left maxilla of the holotype of *Liaoceratops* (IVPP V12738) in distal and labial views; (D), (E) and (F), the cross-sections of A, B and C through the middle part of the roots, showing different outline shape. The arrows indicate where the cross-sections generate. Scale bars equal 10 mm (A-C) and 5 mm (D-F).

*Psittacosaurus lujiatunensis* (IVPP V12617) exhibits similar concave surfaces on the occlusal surface of the crowns as in *Yinlong*, *Chaoyangsaurus,* and *Hualianceratops*. These early-diverging ceratopsians bear similar low-angled wear facets but the depression on the occlusal surface indicates a different occlusion from the shearing occlusal system as in neoceratopsians. In addition, the primary ridges are located at the midline of the crowns in *P.lujiatunensis* (IVPP V12617).

The dental structures in neoceratopsians differ from early-diverging ceratopsians. In *Liaoceratops*, *Archaeoceratops,* and *Auroraceratops*, the crowns developed slightly more prominent and narrow primary ridges and the teeth of *Leptoceratops* and *Protoceratops* developed the most prominent primary ridge outside of ceratopsids (Tanoue et al., 2009). Significantly, the primary ridges in the dentary teeth in *Archaeoceratops* (IVPP V11114) are located relatively mesial to the midline of the crowns in contrast to its maxillary dentitions and other neoceratopsians. Late-diverging neoceratopsians including *Leptoceratops* and *Protoceratops* have deeper indentations mesial and distal to the primary ridge, as in ceratopsids, than early-diverging neoceratopsians (Tanoue et al., 2009). In *Liaoceratops*, *Protoceratops*, *Leptoceratops* and *Zuniceratops* which bear closer-packed dentitions, shallow longitudinal grooves form on the roots to accommodate adjacent crowns in neighboring tooth families. This allows for closer packing of the dentition which may be a precursor of ceratopsid bifid roots (Fig. 9C and 9F) (Brown & Schlaikjer, 1940; Wolfe et al., 1998; He et al., 2018). Among all specimens we examined here, the occlusal surfaces of the functional teeth are regular and generally on the same plane whereas they are irregular in *Protoceratops* and Ceratopsidae (Edmund, 1960; Tanoue et al., 2009; Mallon et al., 2016). Differing from early-diverging ceratopsians, ceratopsids have evolved unique dental features including two-rooted teeth, high angle of the wear facets, a very prominent primary ridge flanked by deep indentations (Edmund, 1960; Tanoue et al., 2009).

#### Replacement progression

The replacement progression in *Yinlong* and *Chaoyangsaurus* differs slightly from *Liaoceratops* (He et al., 2018). The resorption of the functional tooth in *Liaoceratops* is initiated after the successional tooth grew, in contrast to *Yinlong* and *Chaoyangsaurus* (He et al., 2018). When the replacement tooth growth is nearly complete, the labial dentine of the roots in *Liaoceratops* is more completely preserved than in *Yinlong* (Fig. 7C and 7F). In addition, the root of the replacement tooth in *Liaoceratops* inclines lingually at 24° and that in *Yinlong* also inclines lingually but with a smaller angle of inclination (12°), while the root of the replacement tooth in *Chaoyangsaurus* is relatively vertical and is appressed to the functional tooth (Fig. 7 A-C). As a result, the far labial side of the root in *Liaoceratops* and *Yinlong* possibly lies beyond the zone of resorption and the dentine of the functional tooth next to the replacement tooth is still preserved, while that in *Chaoyangsaurus* is resorbed (Fig. 7) (He et al., 2018). In general, the degree of resorption of the functional tooth root is most severe in *Chaoyangsaurus* followed by *Yinlong,* and it is the weakest in *Liaoceratops*. In addition, the functional crown detaches from the root in *Liaoceratops* and the functional root remnants are still present labial to the replacement tooth while the functional tooth is shed (He et al., 2018). The relatively slight resorption and the separation between the resorbed functional crown and root may interpret why the remnants of the functional teeth are so prevalent in *Liaoceratops*.

At present, the replacement process in ceratopsids has not been described in detail. Some transverse sections previously reported suggested a difference in the replacement process between ceratopsids and early-diverging ceratopsians (Erickson et al., 2015). In ceratopsids, the replacement teeth germinated inside the pulp cavities of the predecessors instead of lingual to the predecessors (Erickson et al., 2015 Fig. 1B). The transition of the location of the replacement teeth germinated from the lingual side of the functional roots to the tip of that has been reported in *Leptoceratops* (Brown & Schlaikjer, 1940), which may be a primitive feature of ceratopsids. As such the roots of the preceding functional teeth became two-rooted in ceratopsids (Erickson et al., 2015 Fig. 1B). As the teeth developed, the long axes of the replacement teeth in the same alveolus inclined from labially to lingually (Erickson et al., 2015 Fig. 1B). The roots of the preceding functional teeth in ceratopsids would shed after the crowns have been worn away instead of mostly resorbed as they do in early-diverging ceratopsians (Edmund, 1960).

#### Tooth replacement pattern

Besides the morphological differences, a high rate of tooth replacement characterizes ceratopsids, identified by more replacement teeth in each vertical series (Erickson, 1996). In early-diverging neoceratopsians (*Liaoceratops*, *Auroraceratops*), an alveolus bears at most two replacement teeth with a relatively lower replacement rate (Tanoue et al., 2012; He et al., 2018; Morschhauser et al., 2018). In more early-diverging species of the clade (*Yinlong*, *Chaoyangsaurus*, *Psittacosaurus*, *Hualianceratops*), each alveolus bears at most one replacement tooth indicating lower replacement rates than late-diverging ceratopsians (Table. 3).

**TABLE 3.**
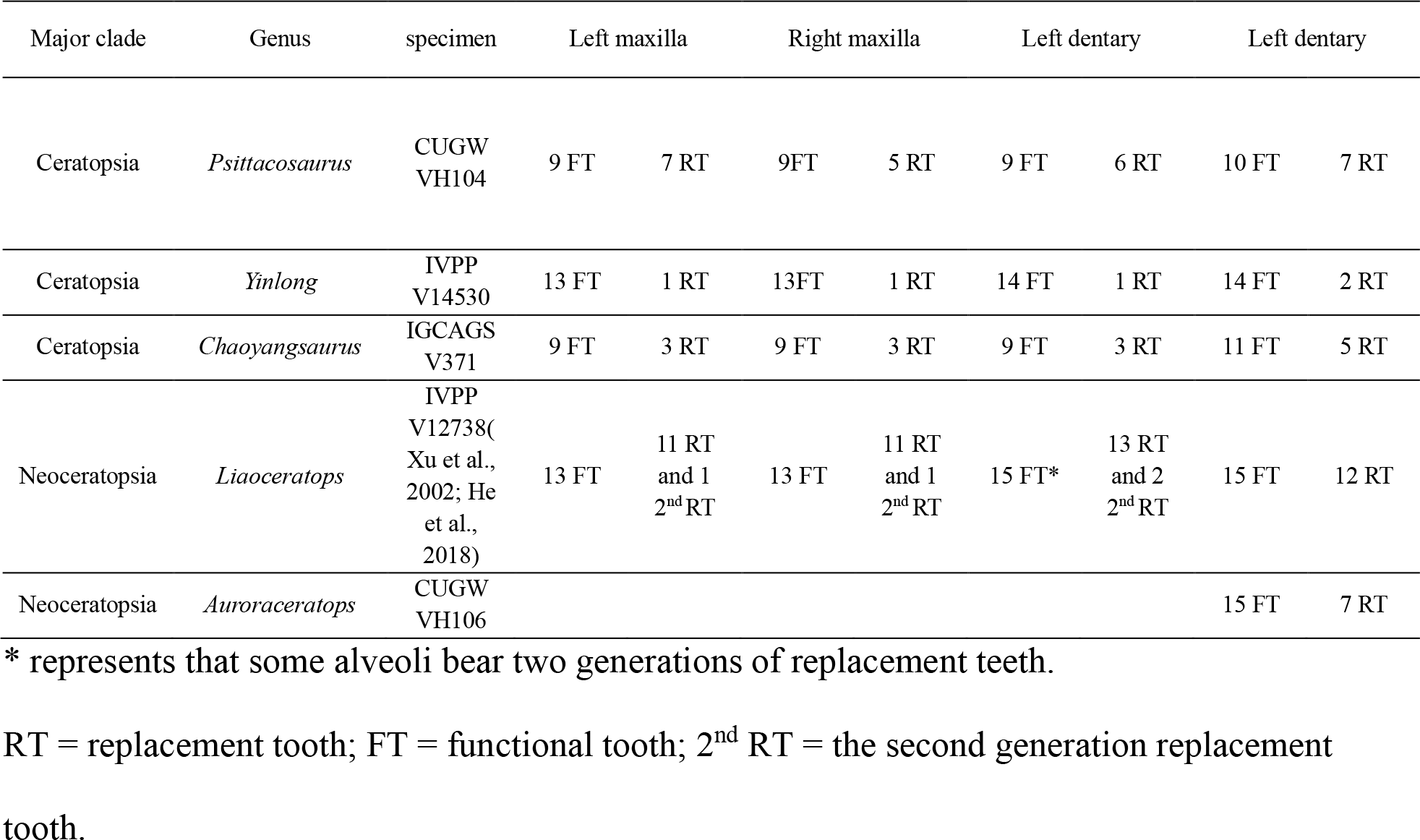
List of the ratio of the replacement teeth to the functional teeth in some ceratopsians which have been studied by computed tomography.

Overall, the evolution of dentitions from the earliest-diverging ceratopsians to ceratopsids are as follows: the development of the primary ridges and the deep indentations; the increased angle of the wear facets on the crowns; the increase of tooth counts in tooth rows; the presence of the shallow grooves on the roots trending from single-rooted teeth to two-rooted teeth; the arrangement of teeth into a more compact mass; the increase of teeth in each tooth family; the location of the replacement teeth transferring from the lingual side of the functional teeth to the inside of the pulp cavities (Fig. 10).

**FIGURE 10.**
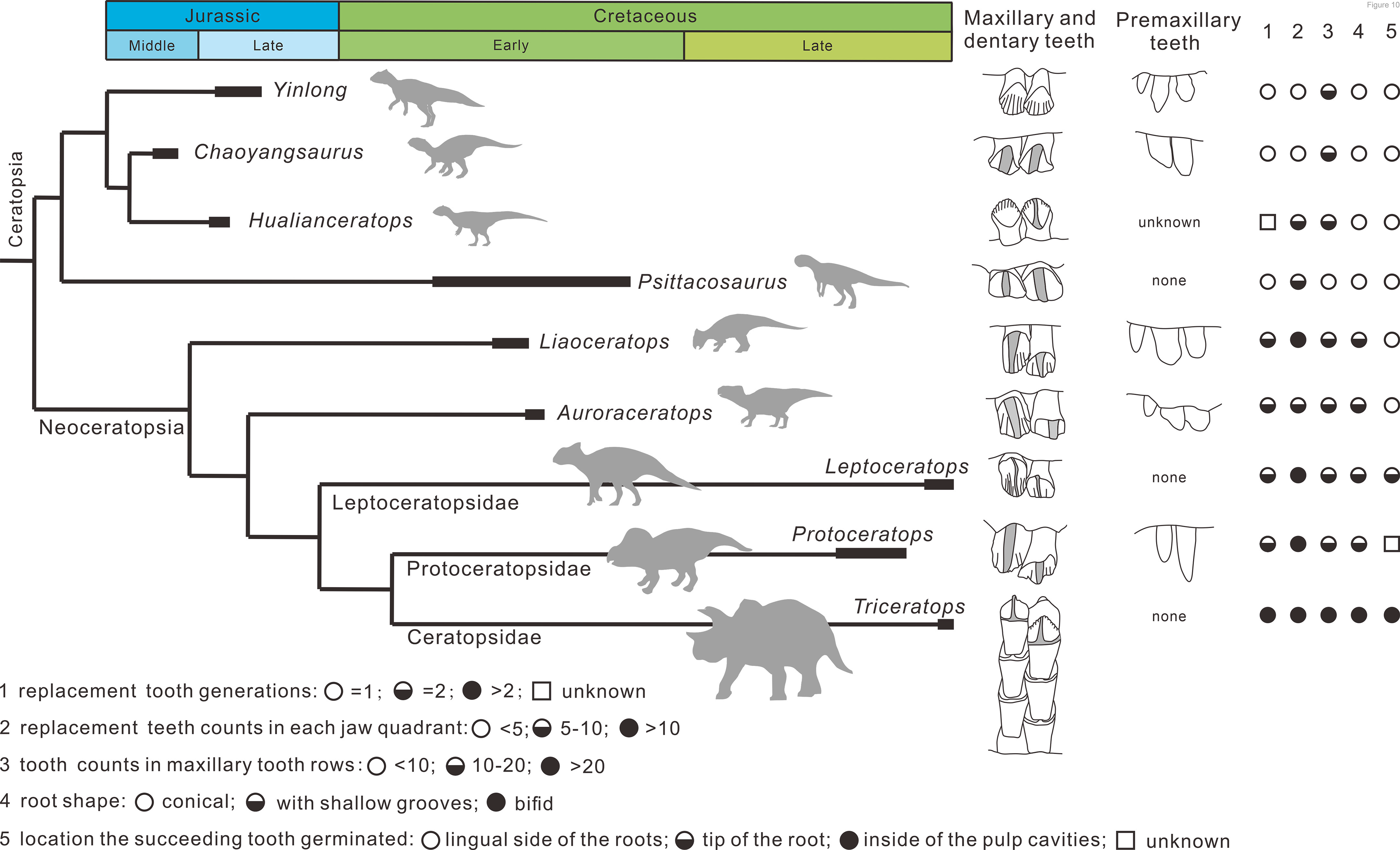
Phylogenetic tree of ceratopsians (composite from Erickson et al. (2015), Han et al. (2018), and Yu et al. (2020)) and comparison of the dental anatomy and the tooth replacement pattern. *Psittacosaurus* from Averianov et al. (2006); *Liaoceratops* from He et al. (2018); *Auroraceratops* from Tanoue et al. (2012) and Morschhauser et al. (2018); *Leptoceratops* from Tanoue et al. (2009); *Protoceratops* from Edmund (1960) and Brown & Schlaikjer (1940); *Triceratops* from Edmund (1960).

### Implication of diet and environment

The upper half of the Shishugou Formation, in which the bonebeds containing *Yinlong* and *Hualianceratops* occur, indicate a warm and seasonally dry climate in the Middle and Late Jurassic (Eberth et al., 2001; Clark et al., 2004; Eberth et al., 2006; Bian et al., 2010; Eberth et al., 2010). Ashraf et al. (2010) thought that the dry season became more and more dominant during the period. Wang et al. (2000) have described the megaplant fossils and identified pollen and spores of some ferns from the Upper Jurassic, Junggar Basin, Xinjiang, China. Based on these identified plants, a high-resolution digital computer model has been reconstructed (Hinz et al., 2010). During the Late Jurassic, this area developed forests near the banks of rivers under moist conditions and consisted primarily of conifers like Araucariaceae and the understory of the forest mainly consisted of *Angiopteris*, *Osmunda* and *Coniopteris* (Mcknight et al., 1990; Hinz et al., 2010). Maiorino et al. (2018) pointed out that *Yinlong* was not able to tolerate high loadings due to its more primitive lower jaw morphology, and they may have fed on softer foliage and fruits or swallowed the food in a relatively unprocessed form. The reconstruction of the flora does not support enough soft food for *Yinlong*. Therefore, we suggest that *Osmunda regalis* as a representative of the fern family Osmundaceae can grow up to 1.5 m in height which is suitable for *Yinlong downsi* to feed (Xu et al., 2006; Hinz et al., 2010). Ferns high in fiber and sand ingestion brought by dry climate likely increase stress in teeth.

Although the tooth replacement rate in *Yinlong* is not clear, previous researches have suggested that the tooth replacement rates in some sauropods and hadrosaurids, which have elaborate dental batteries, are relatively fast (D’Emic et al., 2013). The low number of replacement teeth in *Yinlong* likely reflects slow tooth replacement rates which would not imply rapid tooth wear. To maintain the function of dentitions, *Yinlong* possibly has feeding strategies other than only grinding food with their dentitions. Combined with the low degree of wear in maxillary and dentary teeth, food is possibly simply chewed and then ingested and processed in the hind-gut. Xu et al. (2006) mentioned that seven gastroliths were found within the ribcage of IVPP V14530. These gastroliths are generally accepted to assist food trituration and may have the function to help *Yinlong* to consume highly fibrous plant material (Wings, 2007). Previously, unambiguous evidence supporting the presence of lithophagia is known from *Psittacosaurus* among ceratopsians (Osborn, 1923; Ignacio, 2008). Therefore, some early-diverging ceratopsians that show relatively slow tooth replacement rates and lack evidence of heavy tooth wear likely used gastroliths to triturate foodstuffs to cope with the stringent requirements for digestion of plant materials.

Several morphological adaptations occurred during the evolution of Ceratopsia including the longitudinal ridge of ceratopsids and thickening of the lower jaw in early-diverging neoceratopsians besides the transition of dentitions mentioned above (Bell et al., 2009; Maiorino et al., 2018). Finite element analysis on the lower jaws of ceratopsians suggests that ceratopsids represent the clade with the most efficient masticatory apparatus in Ceratopsia whereas the early- diverging ceratopsians *Hualianceratops* and *Yinlong* had a primitive lower jaw (Maiorino et al., 2018). These changes undoubtedly improve the chewing ability in neoceratopsians and ceratopsids. Given their body difference, the greater food consumption brought by the increased body size may have driven, in part, the evolution of the jaw and the replacement patterns.

However, increased body size may not be the only reason for increased replacement tooth number and the stronger jaw. *Liaoceratops* and *Psittacosaurus* are similar in size to *Yinlong* but have more replacement teeth than *Yinlong* as well as two generations of replacement teeth in Liaoceratops and the jaws able to withstand higher stress (He et al., 2018; Maiorino et al., 2018). The Jehol flora, which occurs in the Yixian Formation of Liaoning, is dominated by Cycadopsida and Coniferopsida (Deng et al., 2012). It suggests that *Liaoceratops* had a different diet strategy from *Yinlong*. Likewise, one of the greatest changes in terrestrial ecosystems during the Cretaceous period was the diversification of angiosperms (Barrett & Willis, 2001). Changes in the floral composition may have resulted in the different diet strategies in ceratopsids. Therefore, the different diet strategies may help explain the different replacement patterns.

## Acknowledgements

We thank the members of the Sino-American expedition team for collecting the fossils described herein, and L. S. Xiang, T. Yu, and X. Q. Ding for preparing the fossils. Yun Feng and YM. Luo for helping CT scan. Yang Wu and Yuzheng Ke for helping reconstruct CT models. This project was supported by the National Natural Science Foundation of China (41972021, 41688103) and the International Partnership Program of Chinese Academy of Sciences (132311KYSB20180016).

## References

1. Ashraf AR, Sun YW, Sun G, Uhl D, Mosbrugger V, Li J, and Herrmann M. 2010. Triassic and Jurassic palaeoclimate development in the Junggar Basin, Xinjiang, Northwest China—a review and additional lithological data. Palaeobiodiversity and Palaeoenvironments 90:187–201. 10.1007/s12549-010-0034-0

2. Averianov AO, Voronkevich AV, Leshchinskiy SV, and Fayngertz AV. 2006. A Ceratopsian dinosaur Psittacosaurus sibiricus from the Early Cretaceous of West Siberia, Russia and its phylogenetic relationships. Journal of Systematic Palaeontology 4:359–395. 10.1017/S1477201906001933

3. Barrett PM, and Willis KJ. 2001. Did dinosaurs invent flowers? Dinosaur–angiosperm coevolution revisited. Biological Reviews 76:411–447. 10.1017/S1464793101005735

4. Becerra MG, Pol D, Whitlock JA, and Porro LB. 2020. Tooth replacement in *Manidens condorensis*: baseline study to address the replacement pattern in dentitions of early ornithischians. Papers in Palaeontology 7:1167–1193. 10.1002/spp2.1337

5. Bell PR, Snively E, and Shychoski L. 2009. A Comparison of the Jaw Mechanics in Hadrosaurid and Ceratopsid Dinosaurs Using Finite Element Analysis. The Anatomical Record 292:1338–1351. https://doi.org/10.1002/ar.20978

6. Bian WH, Hornung J, Liu ZH, Wang P, and Hinderer M. 2010. Sedimentary and palaeoenvironmental evolution of the Junggar Basin, Xinjiang, Northwest China. Palaeobiodiversity and Palaeoenvironments 90:175–186. 10.1007/s12549-010-0038-9

7. Bramble K, LeBlanc ARH, Lamoureux DO, Wosik M, and Currie PJ. 2017. Histological evidence for a dynamic dental battery in hadrosaurid dinosaurs. Sci Rep 7:15787. 10.1038/s41598-017-16056-3

8. Brown DB, and Schlaikjer DEM. 1940. The structure and relationships of *Protoceratops*. Transactions of the New York Academy of Sciences 2:99–100.

9. Chatterjee S, and Zheng Z. 2002. Cranial anatomy of *Shunosaurus*, a basal sauropod dinosaur from the Middle Jurassic of China. Zoological Journal of the Linnean Society 136:145–169. 10.1046/j.1096-3642.2002.00037.x

10. Clark J, Xu X, Forster C, and Wang Y. 2004. New discoveries from the Middle-to-Upper Jurassic Shishugou Formation, Xinjiang, China. Journal of Vertebrate Paleontology Supplement to 24:46A. 10.1016/j.palaeo.2004.06.002

11. Czepiński Ł. 2020. Ontogeny and variation of a protoceratopsid dinosaur Bagaceratops rozhdestvenskyi from the Late Cretaceous of the Gobi Desert. Historical Biology 32:1394–1421. 10.1080/08912963.2019.1593404

12. D’Emic MD, Whitlock JA, Smith KM, Fisher DC, and Wilson JA. 2013. Evolution of high tooth replacement rates in sauropod dinosaurs. Plos One 8:e69235. 10.1371/journal.pone.0069235

13. Demar R. 1972. Evolutionary Implications of Zahnreihen. Evolution 26:435–450.

14. Demar R, and Bolt JR. 1981. Dentitional Organization and Function in a Triassic Reptile. Journal of Paleontology 55:967–984.

15. Deng SH, Lu YZ, Fan R, Li X, Fang LH, and Liu L. 2012. Cretaceous floras and biostratigraphy of Chian. Journal of Stratigraphy 36:241–265.

16. Dodson P, Froster CA, and Sampson SD. 2004. Ceratopsidae. In: Weishampel DB, Osmólska H, and Dodson P, eds. The Dinosauria. Second edn ed. Berkeley: University of California Press, 861.

17. Eberth D, Xu X, Clark J, Machlus M, and Hemming S. 2006. The dinosaur-bearing Shishugou Formation (Jurassic, northwest China) revealed. Journal of Vertebrate Paleontology 26:58A-58A.

18. Eberth DA, Brinkman DB, Chen PJ, Yuan FT, Wu SZ, Li G, and Cheng XS. 2001. Sequence stratigraphy, paleoclimate patterns, and vertebrate fossil preservation in Jurassic Cretaceous strata of the Junggar Basin, Xinjiang Autonomous Region, People’s Republic of China. Canadian Journal of Earth Sciences 38:1627–1644. 10.1139/e01-067

19. Eberth DA, Xu X, and Clark JM. 2010. Dinosaur death pits from the Jurassic of China. PALAIOS 25:112–125. 10.2110/palo.2009.p09-028r

20. Edmund AG. 1960. *Tooth replacement phenomena in lower vertebrates*: Life Sciences Division, Royal Ontario Museum.

21. Erickson GM. 1996. Incremental lines of von Ebner in dinosaurs and the assessment of tooth replacement rates using growth line counts. Proceedings of the National Academy of Sciences 93:14623–14627. 10.1073/pnas.93.25.14623

22. Erickson GM, Sidebottom MA, Kay DI, Turner KT, Ip N, Norell MA, Sawyer WG, and Krick BA. 2015. Wear biomechanics in the slicing dentition of the giant horned dinosaur *Triceratops*. Science Advances 1:e1500055. 10.1126/sciadv.1500055

23. Fastnacht M. 2008. Tooth replacement pattern of *Coloborhynchus robustus* (Pterosauria) from the Lower Cretaceous of Brazil. Journal of Morphology 269:332–348. 10.1002/jmor.10591

24. Han FL, Forster CA, Clark JM, and Xu X. 2015. A new taxon of basal Ceratopsian from China and the early evolution of Ceratopsia. Plos One 11:e0143369. 10.1371/journal.pone.0143369

25. Han FL, Forster CA, Clark JM, and Xu X. 2016. Cranial anatomy of *Yinlong downsi* (Ornithischia: Ceratopsia) from the Upper Jurassic Shishugou Formation of Xinjiang, China. Journal of Vertebrate Paleontology 36:e1029579. 10.1080/02724634.2015.1029579

26. Han FL, Forster CA, Xu X, and Clark JM. 2018. Postcranial anatomy of *Yinlong downsi* (Dinosauria: Ceratopsia) from the Upper Jurassic Shishugou Formation of China and the phylogeny of basal ornithischians. Journal of Systematic Palaeontology 16:1159–1187. 10.1080/14772019.2017.1369185

27. Hanai T, and Tsuihiji T. 2019. Description of Tooth Ontogeny and Replacement Patterns in a Juvenile *Tarbosaurus bataar* (Dinosauria: Theropoda) Using CT-Scan Data. The Anatomical Record 302:1210–1225. 10.1002/ar.24014

28. He YM, Makovicky PJ, Xu X, and You HL. 2018. High-resolution computed tomographic analysis of tooth replacement pattern of the basal neoceratopsian *Liaoceratops yanzigouensis* informs ceratopsian dental evolution. Sci Rep 8:5870. 10.1038/s41598-018-24283-5

29. Hinz JK, Smith I, Pfretzschner HU, Wings O, and Sun G. 2010. A high-resolution three-dimensional reconstruction of a fossil forest (Upper Jurassic Shishugou Formation, Junggar Basin, Northwest China). Palaeobiodiversity and Palaeoenvironments 90:215–240. 10.1007/s12549-010-0036-y

30. Ignacio AC. 2008. Gastroliths in an Ornithopod dinosaur. Acta Palaeontologica Polonica 53:351–355. 10.4202/app.2008.0213

31. Leblanc ARH, Brink KS, Cullen TM, and Reisz RR. 2017. Evolutionary implications of tooth attachment versus tooth implantation: A case study using dinosaur, crocodilian, and mammal teeth. Journal of Vertebrate Paleontology 37:e1354006. 10.1080/02724634.2017.1354006

32. Maiorino L, Farke AA, Kotsakis T, Raia P, and Piras P. 2018. Who is the most stressed? Morphological disparity and mechanical behavior of the feeding apparatus of ceratopsian dinosaurs (Ornithischia, Marginocephalia). Cretaceous Research 84:483–500. 10.1016/j.cretres.2017.11.012

33. Mallon JC, Ott CJ, Larson PL, Iuliano EM, and Evans DC. 2016. *Spiclypeus shipporum* gen. et sp. nov., a boldly audacious new chasmosaurine ceratopsid (Dinosauria: Ornithischia) from the Judith River Formation (Upper Cretaceous: Campanian) of Montana, USA. Plos One 11:e0154218. 10.1371/journal.pone.0154218

34. Mcknight CL, Graham SA, Carroll AR, Gan Q, Dilcher DL, Zhao M, and Liang YH. 1990. Fluvial sedimentology of an Upper Jurassic petrified forest assemblage, Shishu Formation, Junggar Basin, Xinjiang, China. Palaeogeography Palaeoclimatology Palaeoecology 79:1–9. 10.1016/0031-0182(90)90102-D

35. Morschhauser EM, Li DQ, You HL, and Dodson P. 2018. Cranial anatomy of the basal neoceratopsian *Auroraceratops rugosus* (Ornithischia: Ceratopsia) from the Yujingzi Basin, Gansu Province, China. Journal of Vertebrate Paleontology 38:36–68. 10.1080/02724634.2017.1399136

36. Osborn HF. 1923. Two Lower Cretaceous dinosaurs of Mongolia. American Museum Novitates 95:1–10.

37. Ryan MJ, Chinnery-Allgeier BJ, Eberth DA, Ralrick PE, and Currie PJ. 2010. New Perspectives on Horned Dinosaurs: The Royal Tyrrell Museum Ceratopsian Symposium. Bloomington: Indiana University Press.

38. Sereno PC. 2006. Psittacosauridae. In: Weishampel DB, Osmólska H, and Dodson P, eds. The Dinosauria. First edn ed. Berkeley: University of California Press, 861.

39. Sereno PC. 2012. Taxonomy, morphology, masticatory function and phylogeny of heterodontosaurid dinosaurs. ZooKeys:1–225. 10.3897/zookeys.223.2840

40. Tanoue K, Li Dq, and You Hl. 2012. Tooth replacement pattern in maxillary dentition of basal Neoceratopsia. Bulletin of the Kitakyushu Museum of Natural History and Human History Series A (Natural History) 10:123–127. 10.34522/kmnh.10.0_123

41. Tanoue K, You HL, and Dodson P. 2009. Comparative anatomy of selected basal ceratopsian dentitions. Canadian Journal of Earth Sciences 46:425–439. 10.1139/E09-030

42. Wang YD, Zhang W, and Saiki K. 2000. Fossil woods from the Upper Jurassic of Qitai, Junggar Basin, Xinjiang, China. Acta Palaeontologica Sinica 39:176–185.

43. Weishampel DB, Dodson P, and Osmólska H. 2004. The Dinosauria. Berkeley: University of California Press.

44. Wiersma K, and Sander PM. 2017. The dentition of a well-preserved specimen of *Camarasaurus* sp.: implications for function, tooth replacement, soft part reconstruction, and food intake. PalZ 91:145–161. 10.1007/s12542-016-0332-6

45. Wings O. 2007. A review of gastrolith function with implications for fossil vertebrates and a revised classification. Acta Palaeontologica Polonica 52:1–16. 10.1038/sj.onc.1207250

46. Wolfe DG, Kirkland JI, and Lucas SG. 1998. *Zuniceratops christopheri* n. gen. & n. sp., a ceratopsian dinosaur from the Moreno Hill Formation (Cretaceous, Turonian) of west-central New Mexico. Lower and Middle Cretaceous Terrestrial Ecosystems New Mexico Museum of Natural History and Science Bulletin 14:303–317.

47. Xu X, Forster CA, Clark Jm, and Mo J. 2006. A basal ceratopsian with transitional features from the Late Jurassic of Northwestern China. Proceedings of the Royal Society B: Biological Sciences 273:2135–2140. 10.1098/rspb.2006.3566

48. Xu X, Makovicky Pj, Xl X, Norell MA, and You H. 2002. A Ceratopsian dinosaur from China and the early evolution of Ceratopsia. Nature 416:314–317. 10.1038/416314a

49. Yu C, Prieto-Marquez A, Chinzorig T, Badamkhatan Z, and Norell M. 2020. A neoceratopsian dinosaur from the early Cretaceous of Mongolia and the early evolution of ceratopsia. Communications Biology 3:499. 10.1038/s42003-020-01222-7

50. Zhao X, Cheng Z, and Xu X. 1999. The earliest ceratopsian from the Tuchengzi Formation of Liaoning, China. Journal of Vertebrate Paleontology 19:681–691. 10.1080/02724634.1999.10011181

